# A statistical framework for neuroimaging data analysis based on mutual information estimated via a Gaussian copula

**DOI:** 10.1101/043745

**Authors:** Robin A. A. Ince, Bruno L. Giordano, Christoph Kayser, Guillaume A. Rousselet, Joachim Gross, Philippe G. Schyns

## Abstract

We begin by reviewing the statistical framework of information theory as applicable to neuroimaging data analysis. A major factor hindering wider adoption of this framework in neuroimaging is the difficulty of estimating information theoretic quantities in practice. We present a novel estimation technique that combines the statistical theory of copulas with the closed form solution for the entropy of Gaussian variables. This results in a general, computationally efficient, flexible, and robust multivariate statistical framework that provides effect sizes on a common meaningful scale, allows for unified treatment of discrete, continuous, uni-and multi-dimensional variables, and enables direct comparisons of representations from behavioral and brain responses across any recording modality. We validate the use of this estimate as a statistical test within a neuroimaging context, considering both discrete stimulus classes and continuous stimulus features. We also present examples of analyses facilitated by these developments, including application of multivariate analyses to MEG planar magnetic field gradients, and pairwise temporal interactions in evoked EEG responses. We show the benefit of considering the instantaneous temporal derivative together with the raw values of M/EEG signals as a multivariate response, how we can separately quantify modulations of amplitude and direction for vector quantities, and how we can measure the emergence of novel information over time in evoked responses. Open-source Matlab and Python code implementing the new methods accompanies this article.

**Highlights:**

- Novel estimator for mutual information and other information theoretic quantities
- Provides general, efficient, flexible and robust multivariate statistical framework
- Validated statistical performance on EEG and MEG data
- Applications to spectral power and phase, 2D magnetic field gradients, temporal derivatives
- Interaction information relates information content in different responses

## 1. Introduction

Mutual Information (MI) measures the statistical dependence between two random variables (Cover and Thomas, 1991; Shannon, 1948). It can be viewed as a statistical test against a null hypothesis that two variables are statistically independent, but in addition its effect size (measured in bits) has a number of useful properties and interpretations (Kinney and Atwal, 2014).

There is a long history of applications of MI for the study of neural activity (Borst and Theunissen, 1999; Eckhorn and Pöpel, 1974; Fairhall et al., 2012; Nelken and Chechik, 2007; Rolls and Treves, 2011; Victor, 2006). MI has been used to compare different neural response codes (Ince et al., 2013; Kayser et al., 2009; Reich et al., 2001), characterization of different neurons (Sharpee, 2014) as well as quantification of the effect of correlations between neurons (Ince et al., 2009, 2010; Moreno-Bote et al., 2014) and of the importance of spike timing (Kayser et al., 2010; Nemenman et al., 2008; Panzeri et al., 2001). Recent studies have begun to explore its application to neuroimaging (Afshin-Pour et al., 2011; Caballero-Gaudes et al., 2013; Gross et al., 2013; Guggenmos et al., 2015; Ostwald and Bagshaw, 2011; Panzeri et al., 2008; Salvador et al., 2007; Saproo and Serences, 2010; Schyns et al., 2011; Serences et al., 2009).

Despite its useful properties, a possible reason for why MI has not been more widely adopted, particularly within the neuroimaging community, is the difficulty of accurately estimating MI from limited quantities of experimental data (Steuer et al., 2002). The most common approach involves quantizing the data into a number of bins, and estimating MI over the resulting discrete spaces. However, this method is sensitive to the problem of limited sampling bias, which is particularly acute when considering multi-dimensional responses. Several continuous methods are available (outlined briefly in Section 2), but while these measures are often less sensitive to sampling bias effects, they can be computationally expensive and often require the estimation (or ad-hoc setting) of additional parameters.

Here we present a novel approach to estimating MI with continuous variables. Our method is rank-based, robust and makes no assumptions on the marginal distributions of each variable. It does make an assumption on the form of the relationship between the variables, which results in the estimate being a lower bound to the true MI. It is computationally efficient and statistically powerful when applied within a permutation-based null-hypothesis testing framework. We highlight the benefits resulting from the ability of the estimator to extend to multivariate response spaces, which are often intractable with other methods. This improved multivariate performance allows estimation of quantities such as Conditional Mutual Information (CMI) (Ince et al., 2012), Directed Information (also called transfer entropy) (Ince et al., 2015; Massey, 1990; Schreiber, 2000) as well as measures quantifying pairwise interactions between variables (Chicharro, 2014; Panzeri et al., 2008). We believe these higher-order information-theoretic quantities have the potential to provide transformative new interpretations of neuroimaging data, by providing a unified framework for analyses based on the information content of neural signals (Kriegeskorte and Bandettini, 2007; Naselaris et al., 2011; Schyns et al., 2009). The methods we present enable study of the representation, processing, and communication in the brain of multiple features of the external world (Ince et al., 2015). Furthermore, they also enable the study of the relationships between representation in different signals (Kriegeskorte et al., 2008). Overall, this new estimator provides the basis for a useful and flexible multivariate statistical framework for the analysis of neuroimaging data.

In this paper, we first introduce the concepts of entropy and MI within a neuroimaging context, and briefly review current MI estimation methods. We also describe higher-order information theoretic quantities and possible neuroimaging applications (Section 2). We then present our novel MI estimator (Section 3) and demonstrate its statistical performance when combined with a permutation-based null-hypothesis testing framework, with simulations and examples on several data sets (Section 4).

## 2. Review of information theory for neuroimaging

**In this section we review information theoretic methods from a neuroimaging perspective. Readers familiar with information theory can skip this section and proceed to Section 3 where we present our novel MI estimator.**

### 2.1. Entropy

Entropy is the foundational quantity of information theory, and is a measure of the uncertainty, or variability, of a random variable. For any particular value of a random variable, a low probability means that outcome is less likely to occur, and so an observer would be more surprised to see it. A high probability value would be less surprising. This notion can be formulated mathematically: the surprise for value *x* drawn from a distribution *P*(*x*) is defined as −log_2_ *P*(*x*). Entropy is then defined as the expected (average) surprise over the distribution (Figure 1). If an observer draws samples from a distribution, a lower entropy distribution means the observer will be less surprised (or uncertain) about the outcome of any particular sample – i.e. they would be able to make a more accurate guess. Spread out distributions (with high variability) will have high entropy since all potential outcomes have similar probabilities and so the outcome of any particular draw is very uncertain. On the other hand, concentrated distributions (with low variability) will have lower entropy, since some outcomes will have high probability, allowing a reasonable guess to be made about the outcome of any particular draw. Figure 1 shows some examples of entropy of some continuous and discrete variables.

**Figure 1.**
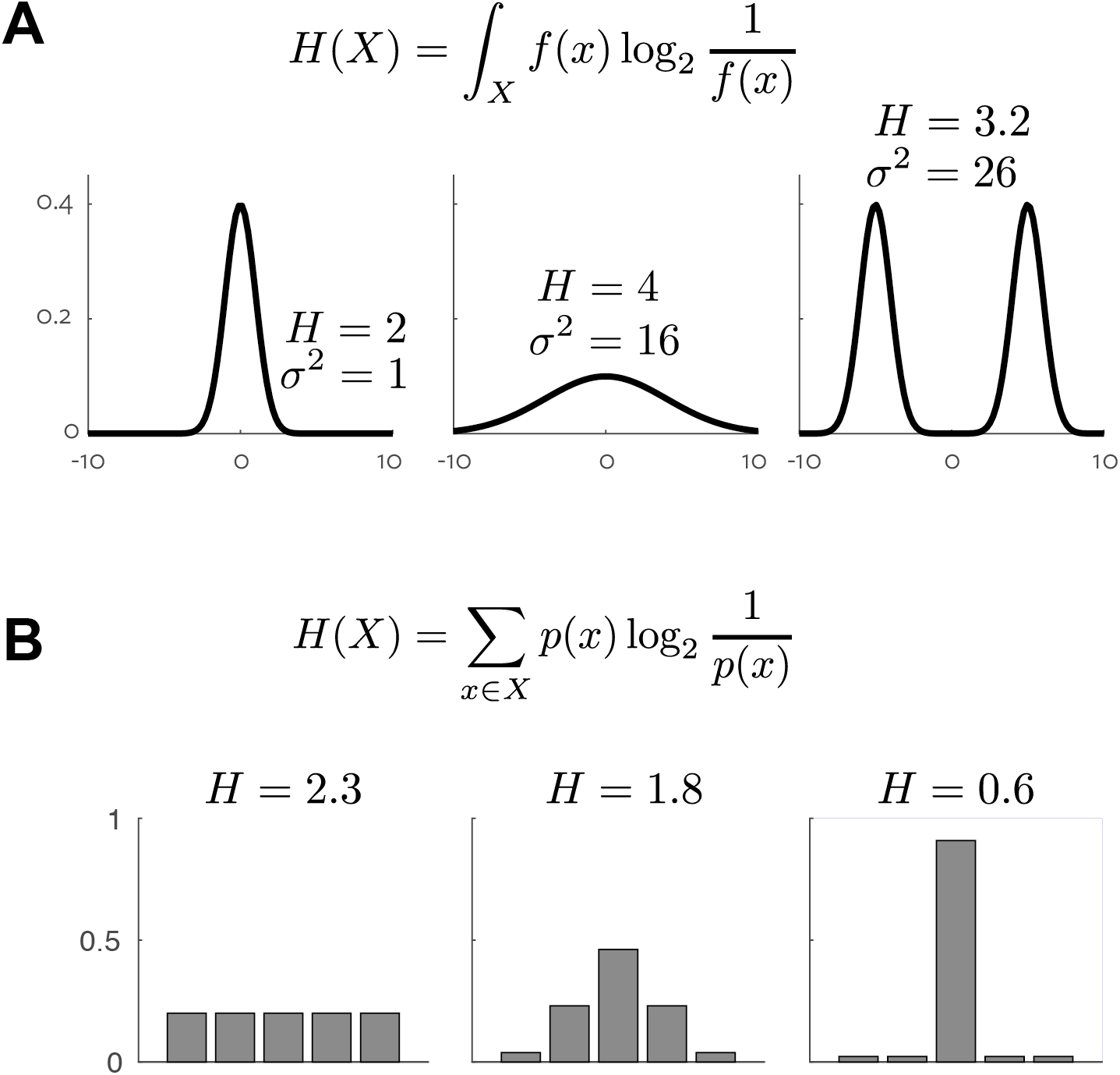
*Entropy values for some example distributions.* **A**. Variance (*σ*^2^) and entropy (*H*) for three example continuous 1D distributions. **B**. Entropy (H) for three example discrete distributions.

When the logarithm used in the definition of surprise is base-2 (Figure 1), the resulting entropy value has units of *bits*. In this case the entropy value has a useful interpretation. If an ideal observer with knowledge of the true distribution has to guess the value of a particular sample by asking a series of yes/no questions, the entropy in bits gives the average number of questions required. Equivalently, a reduction of entropy by 1 bit corresponds to halving of the uncertainty.

If the random variable considered is discrete, the distribution with maximal possible entropy is the uniform distribution (Figure 1B). For continuous valued variables, the term *differential entropy*, is often used – but here for conciseness we use *entropy* for both types of variable. In the continuous case, entropy can be thought of as a generalized form of variance although unlike variance, which is appropriate to use as a measure of spread or dispersion only for unimodal distributions, entropy can give a meaningful quantification of spread for any form of distribution. This analogy with variance can be useful to keep in mind when considering other information theoretic quantities (Garner and McGill, 1956). In the case of variables taking continuous values (i.e. with infinite support), for a specified mean and variance the distribution with maximal entropy is a Gaussian. Further, for Gaussian distributions the entropy is proportional to the logarithm of the variance.

Entropy alone can form a useful measure of the complexity of a signal (Abásolo et al., 2006; Inouye et al., 1991; Overath et al., 2007). However here we are interested primarily its relation to mutual information, which quantifies the relationship between two variables (for example an external stimulus and a recorded signal) in terms of differences in entropies.

### 2.2. Mutual information

Mutual information (MI) is a measure of the statistical dependence between two random variables (Cover and Thomas, 1991; Latham and Roudi, 2009). It is the most general such measure because MI makes no assumptions on the distribution of the variables, or the nature of the relationship between them and is sensitive to non-linear and non-monotonic effects (Figure 2).

**Figure 2.**
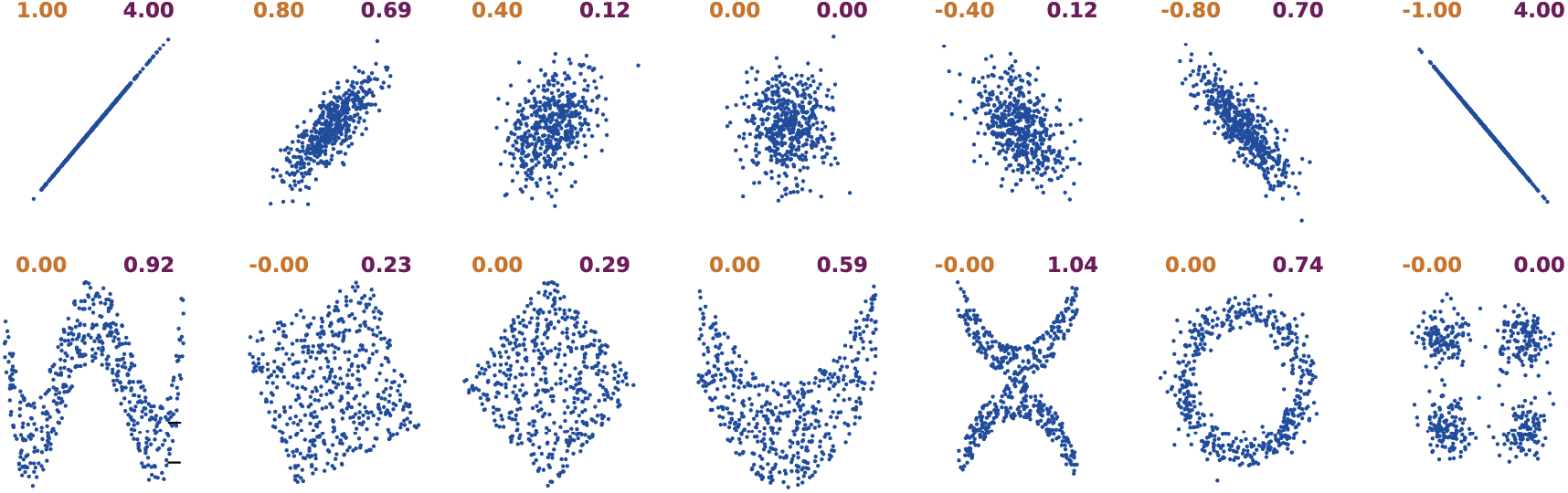
*Examples of correlation vs mutual information.* Each panel illustrates a scatter plot of samples drawn from a particular bi-variate distribution. For each example, the correlation between the two variables is shown in orange (left) and the MI is shown in purple (right; discrete method, 16 bins, 100,000 samples, no bias correction). The top row shows linear relationships, for which MI and correlation both detect a relationship (although on different scales, and note that as MI is always positive it does not reveal the direction of the relationship). The bottom row shows a series of distributions for which the correlation is zero.

MI is defined in terms of entropy differences. As a motivating example, consider a roll of a fair 6-sided die. The outcome of any particular roll follows a uniform distribution over the 6 possible values, with entropy log_2_ 6. If an observer is told that the result of a particular roll is an even number, there are now 3 possible values, all equally likely, and the entropy of this distribution is log_2_ 3. The difference between these entropies is 1 bit:

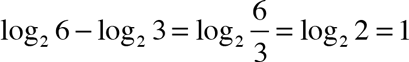

Thus, 1 bit quantifies the amount of information conveyed by the knowledge “this roll is even”, and corresponds to a halving of the uncertainty about the outcome (from 6 possibilities to 3).

There are three mathematically equivalent ways to define MI based on entropy differences, each providing a different perspective on the interpretation of the resulting measure. For illustration within a neuroimaging context, *S* denotes a random variable representing some stimulus feature that is varied across multiple presentations (e.g. in the visual domain, edge contrast, or orientation, or opacity; in the auditory domain loudness, pitch and so forth) and *R* some neural response (e.g. EEG voltage, MEG source amplitude or fMRI bold voxel response measured at a specific site at a specific post-stimulus latency).

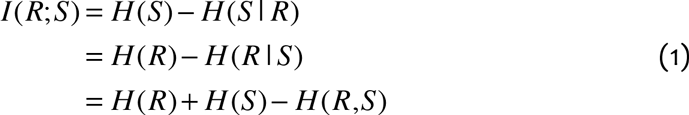

Here, *H* (*S* | *R*) is the conditional entropy: the expectation over values *r* of *R* of the entropy of the distribution of *S* conditional on *r*. *H* (*R*,*S*) represents the entropy of the joint distribution of *R* and *S* (i.e. the two dimensional variable obtained by combining R and S).

The first expression in Eq. (1) shows that MI quantifies the average reduction in uncertainty about which stimulus *S* (e.g. edge contrast, or auditory pitch) was presented after observation of a response *R* (e.g. MEG source amplitude or fMRI bold response). The second expression demonstrates the symmetry of MI, and shows that it equally quantifies the average reduction in uncertainty about the neural response when the stimulus is known. Here it is useful to revisit the analogy with variance – MI measures the *entropy explained* by knowledge of the second variable, which is conceptually similar to the notion of *variance explained* with linear correlation. However, unlike linear correlation, MI makes no assumption on the form of the relationship.

The third expression in Eq. (1) shows that MI quantifies the difference in entropy between a model in which the two variables are statistically independent and the true joint distribution (*H* (*R*,*S*)). The statistically independent model is given by the product of the marginal distributions of the two variables, with entropy *H*(*R*)+*H*(*S*). MI can also be expressed as the Kullback-Leibler (KL) divergence (a measure of distance between probability distributions) between the statistically independent model and the true joint distribution (Akaike, 1992). In fact, in the discrete case, if the probability distributions are estimated via a histogram (multinomial maximum likelihood) method and then used to estimate MI directly from the definition, this estimate is proportional (with a scale factor depending on the number of data points) to the effect size for the log-likelihood ratio test of independence, often called the G-test (Sokal and Rohlf, 1981). The G-test statistic is equal to this maximum likelihood direct MI estimate, multiplied by a factor 2*N* log(2), and is chi-square distributed with the same degrees of freedom as the corresponding chi-square test: 2*N*log(2)*I* ~ χ^2^(*df*), *df* = (|*R*| − 1)(|*S*| − 1). The Neyman-Pearson lemma (Neyman and Pearson, 1933) states that for a given significance level, the likelihood ratio test is the most powerful statistical test for comparing two nested models (used here to test for independence). This motivates perhaps the most useful interpretation of MI from a neuroimaging perspective: a statistical test for independence. It is worth repeating that all three expressions above are mathematically equivalent, but considering them separately explains the different interpretations that can be applied to MI.

MI has several useful properties that are worth highlighting. As discussed above, it is symmetric in the variables considered. It is also additive for independent variables. Additivity derives directly from the mathematical properties of the logarithm: if two variables are statistically independent their joint probability is, by definition, the product of their individual probabilities and therefore the log joint probability is the sum of the individual log probabilities. This means the joint entropy of two independent variables is the sum of the individual entropies. Similarly, the MI between independent pairs of variables is added when they are considered jointly. Formally, two pairs of variable are independent if the full joint probability over all four variables factors as the product of the pairwise joint probabilities:

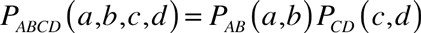

In this case the information conveyed by both pairs is the sum of that conveyed by each pair: *I*(*A*, *C*; *B*, *D*) = *I*(*A*; *B*) + *I*(*C*; *D*). This is a crucial property that is not shared by the effect sizes of other statistical tests and which enables direct quantification of pairwise interaction effects (see Section 2.6). MI measures dependence on a common scale (bits), which provides a meaningful effect size (Friston, 2012) and allows direct comparisons across different responses, experimental modalities or with behavior.

Within the field of information theory there are many mathematical results revealing further properties of the MI measure. A theorem called the Channel Coding Theorem (Cover and Thomas, 1991) relates MI to the transmission capacity of noisy communication channels. A noisy channel can be represented by a conditional probability distribution quantifying the relationship between the output symbols *y* and the input signals *x*: *P_Y_* _|*X*_ (*y* | *x*). This is a fixed property of the channel, but the MI between *x* and *y* depends also on the distribution of the inputs *P_X_* (*x*). The maximum rate at which information can be transmitted over the noisy channel without errors (the channel capacity) is given by the maximal value of MI over all possible input distributions. MI was originally developed within this coding framework, which represents communication as the transmission of a set of discrete symbols over a noisy channel. MI provides theoretical limits on communication efficiency and helped to formulate coding principles that are now pervasive in modern communication systems. This interpretation also motivated much of the early application of MI within neuroscience, viewing the neural pathway from the stimulus receptors to the recorded brain activity as a noisy communication channel, and using MI to quantify properties of this putative communication channel as well as investigating encoding and decoding schemes at different neural levels. Another theorem called the Data Processing Inequality (Cover and Thomas, 1991) states that post-processing cannot increase information. Formally, if the response *R* is transformed to a new representation *P*, where *P* is a probabilistic function of *R* only (and does not depend on *S*), then *I*(*R*; *S*) ≥ *I*(*P*; *S*). This is a desirable property for a neuroimaging statistic as it ensures that any signal processing or feature extraction applied to the recorded responses (for example spectral analysis) cannot artificially inflate the measured effect size, provided it is applied across the whole data set without incorporating knowledge of the stimulus.

Given the above, we suggest there are two views that can be adopted when applying MI in practice (Nelken and Chechik, 2007). The first relies on the coding interpretations of MI, and therefore requires accurate and bias-free estimates; this view has driven most neuroscience applications of MI to date. The second view considers MI more like a conventional statistical hypothesis test of independence, comparable to a t-test or correlation. With this view, accurate bias-free values are less important, but determining statistical significance, including accounting for the problem of multiple comparisons, is crucial. We suggest that this second view is more useful for neuroimaging. In comparison to other statistical tests, MI brings the advantages of sensitivity and robustness (demonstrated in Results), as well as additivity, which allows direct comparisons of the MI from different neural responses. With the novel estimator presented here, MI allows treatment of discrete, continuous and possibly multidimensional variables within a common framework with directly comparable effect sizes on a common and meaningful scale. It should be noted that in practice, as for any statistical test, the measured effect size depends both on the strength of the functional relationship present but also on the noise level of the recorded signal (the signal to noise ratio or SNR). MI, like any statistical test, effectively quantifies the strength of modulation of the signal by a stimulus feature or experimental condition within the particular noise profile of that signal. This should be kept in mind when comparing these effect sizes between different signals. However, the effect sizes from other statistical tests, while also being affected by SNR are also often strongly dependent on parameters such as the number of samples or the particular degrees of freedom in the experimental design making quantitative comparison between results much more difficult.

### 2.3. Existing methods for calculating mutual information

There are several different practical approaches for calculating MI from experimental observations, which we briefly review here from a neuroimaging perspective.

#### Binned methods

Although neuroimaging signals typically take continuous values (voltage in EEG, magnetic field strength in MEG, fMRI BOLD signal amplitude) one commonly used approach for such signals consists of quantizing the continuous valued observations into a set of discrete categories or bins. We therefore briefly review methods for estimating information values on discrete spaces. There are two main strategies for the quantization step: either bins are set with equal spacing, or bins are sized to have approximately equal occupancy (i.e. if using four bins, each data sample is labeled according to the quartile of the empirical distribution in which it lies). For data that are approximately normally distributed, the second method is preferable, as fixed width bins result in the extreme bins having few samples, which exacerbates the limited sampling problem. This is the simplest approach; much work has considered different methods to optimize this quantization step (Darbellay and Vajda, 1999; Endres and Foldiak, 2005; Fraser and Swinney, 1986; Reshef et al., 2011).

Once the continuous signal is quantized, a common approach is to apply multinomial maximum likelihood estimation (the histogram estimate) of the underlying distributions and calculate entropy and MI from their definitions using these distributions (e.g. combining the entropy definition of Figure 1A with Eq (1)). This approach is variously called the naïve, direct or plug-in estimate (Cover and Thomas, 1991). As noted earlier, the Neymann-Pearson lemma states that the likelihood-ratio test (equivalent to MI) is the most powerful hypothesis test for a given significance level (Neyman and Pearson, 1933), a fact which, coupled with the computational efficiency of calculating binned MI, demonstrates the potential usefulness of this quantity as an exploratory statistical test for neuroimaging. Other properties of the MI effect size, such as additivity, provide additional advantages (Section 2.6).

However, when trying to obtain accurate MI estimates for interpretation from a coding channel perspective, there is a problem that the plug-in estimate described above is biased upwards; estimates calculated from finite numbers of samples will be higher than the true value even when averaged over many sample sets. A wide variety of approaches have been proposed to address this problem (Paninski, 2003; Panzeri et al., 2007). The simplest approach is to subtract the mean of the distribution expected under the null hypothesis that there is no relationship between the two variables. As can be seen from the relationship with the chi-square distribution described earlier this is given by (|*R*| − 1)(|*S*| − 1)/2*N* log(2), where N is the number of samples and |*R*|, |*S*| represent the cardinality of the two discrete input spaces. This is called the Miller-Madow correction (Miller, 1955). Various extensions have been proposed to deal with this in different situations such as Bayesian approaches (Nemenman et al., 2004; Panzeri and Treves, 1996), specific methods to estimate entropy and information rates in ongoing processes (Kennel et al., 2005; Shlens et al., 2007) and to deal specifically with the sparse binary probability spaces that result from measuring single neuron spiking activity (Archer et al., 2013; Montemurro et al., 2007). However, we stress again that if the goal of the analysis is classical statistical inference, bias correction is not necessary, and can actually reduce statistical power due to the increased variance of bias corrected estimators (Ince et al., 2012).

A further consideration with binned methods is that they suffer from the curse of dimensionality (Geman et al., 1992). The number of parameters that must be estimated for the multinomial distributions grows exponentially with the number of variables considered. This renders it practically impossible to apply this approach to calculate MI and higher order information-theoretic quantities (Sections 2.8, 2.9) from multivariate spaces with the amounts of data that can be realistically collected from neuroimaging experiments. It may be possible to exploit techniques such as dimensionality reduction or clustering approaches, often referred to as vector quantization in this context (Wilcox and Niles, 1995), to directly quantize multivariate spaces into a small number of representative symbols. However, such an approach also removes the possibility of investigating the effects of the different variables in the multivariate space, for example by considering the MI of each variable individually, or investigating the effect of correlations between them (Chicharro, 2014; Magri et al., 2009; Panzeri and Treves, 1996).

#### Continuous methods

Several methods exist for estimating MI between continuous variables without the quantization step. The most direct way is to first estimate the continuous probability distributions with a Kernel Density Estimation (KDE) technique, and then numerically integrate those estimates to obtain MI (Moon et al., 1995). An alternative approach, which bypasses explicit estimation of the joint distributions, exploits the relationship between probability density and local neighborhood structure. These methods estimate entropy and MI using k-nearest neighbor structure (Faivishevsky and Goldberger, 2009; Kraskov et al., 2004; Victor, 2002) – the probability densities are estimated implicitly from the pairwise distances between samples. These methods have also been extended through careful choice of the distance metric used. For example, for spike trains, various metrics can be defined that emphasize different properties of the spike trains (Victor, 2005). MI conveyed by high-dimensional time courses can be estimated based on a hyperbolic distance measure formed from the correlation coefficient between pairs of time series (Afshin-Pour et al., 2011). While these methods are relatively unbiased, they often have a high variance and are computationally intensive. This is particularly problematic when combined with permutation testing in a mass-univariate neuroimaging context (Groppe et al., 2011).

An alternative approach to dealing with continuous data is to assume a parametric form for the distribution. For example, Local Field Potentials (and similarly M/EEG data) are often approximately Gaussian (Magri et al., 2009). The parameters of this assumed distribution can be estimated from the data and the entropy and MI values estimated directly from the parametric model, for which there are often closed form expressions solving the integral definition of these quantities. While different parametric models can be used (for example, a t-distribution might be more appropriate for M/EEG data), and this approach is computationally efficient, it is not clear what effect a violation of the parametric assumptions would have on the estimate. Further, there are many variables of interest, for example stimulus features obtained from dynamic naturalistic stimuli, which do not have an obviously appropriate parametric form.

### 2.4. Estimating the entropy and MI of Gaussian variables

For Gaussian variables, the integral definition (Figure 1A) can be solved analytically resulting in a closed form expression for the entropy (in bits) as a function of the determinant of the covariance matrix Σ (with dimensionality *k*):

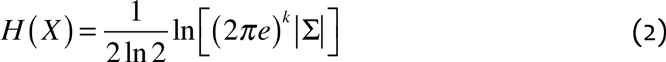

This measure still exhibits some bias due to the estimation of the covariance matrix from limited data, but there exists an analytic correction to remove much of this effect (Goodman, 1963; Magri et al., 2009; Misra et al., 2005). The bias-corrected entropy estimate is given by:

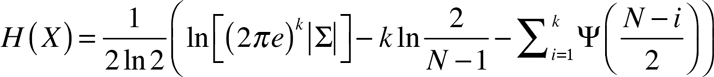

where *N* is the number of samples and *k* is the dimensionality of X with covariance matrix Σ.

From equation (1), the MI between two Gaussian variables is therefore given by:

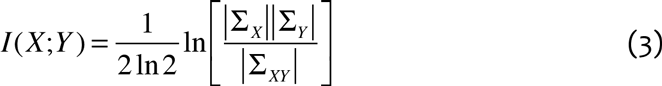

where Σ_*X*_, Σ_*Y*_ are the covariance matrices of X and Y respectively and Σ_*XY*_ is the covariance matrix for the joint variable (*X*,*Y*).

Figure 3 shows the MI between two 1-D Gaussian variables as a function of their correlation. This reveals two key properties. First, the symmetric shape of the graph demonstrates how, since MI is an unsigned quantity, it can reveal the strength but not the direction of a relationship; this is an important aspect to keep in mind. Second, the relationship is clearly non-linear. We suggest that this non-linearity is an advantage, especially for neuroimaging studies, since it results in an enhanced contrast of strong effects with respect to background values in mass-univariate analyses.

**Figure 3.**
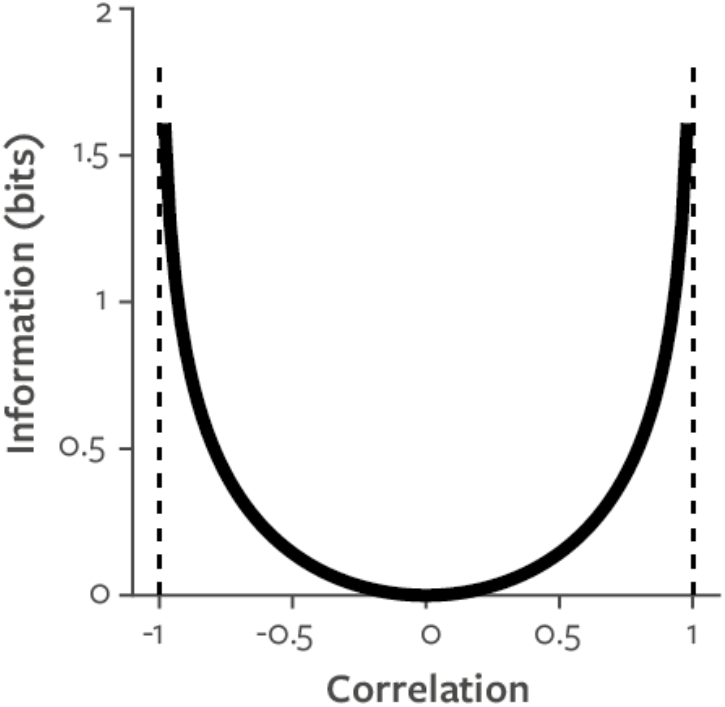
*Relationship between correlation and information for two 1-D Gaussian variables.*

While the parametric Gaussian approach is data robust due to the relatively low number of parameters that need to be estimated, it is not clear how the estimator might perform if the Gaussian distribution assumption was violated, and it cannot be employed in many cases where the distribution of stimulus values is highly non-Gaussian.

### 2.5. Estimating MI within different types of experimental design

MI is, like Pearson correlation, a function of two variables that can be applied in practice to many different sorts of data. In order to correctly interpret a particular information theoretic analysis it is important to understand how the samples used to estimate MI were obtained. Given the diversity of experimental designs employed in neuroimaging, the different approaches to obtaining samples are often a point of confusion. In this section we describe some common experimental designs and detail how MI is estimated in each case.

#### Event related design

An event-related design consists of serial presentation of stimuli, possibly from different classes or with parametrically varying features, and possibly requiring a behavioral response. These presentations are separated in time and the analysis begins by extracting sections of neuroimaging recordings following each presentation. Event-related experiments are typically analyzed by averaging response epochs to different stimulus classes (resulting in an Event-Related Potential or ERP) and performing group statistics. However, using MI we can also quantify modulation of the M/EEG response by a continuous stimulus feature (e.g. a variation of stimulus orientation) that varies across trials. To apply MI in this paradigm, each post-stimulus time point and sensor/source is treated independently within a mass-univariate framework (Groppe et al., 2011). The MI calculation is repeated for each time point and sensor/source, using the repeated presentations of the stimulus as samples (Figure 4A). Multiple comparison correction is required over time points and sensors/sources: this can be achieved using permutation testing (repeating the calculation with shuffled stimulus values) combined with the method of maximum statistics (Holmes et al., 1996; Nichols and Holmes, 2002), or cluster sum statistics (Maris and Oostenveld, 2007) possibly with threshold-free cluster enhancement (Pernet et al., 2015; Smith and Nichols, 2009). An advantage of this design is that, because each time point is analyzed separately, there is no assumption that the signal is stationary.

**Figure 4.**
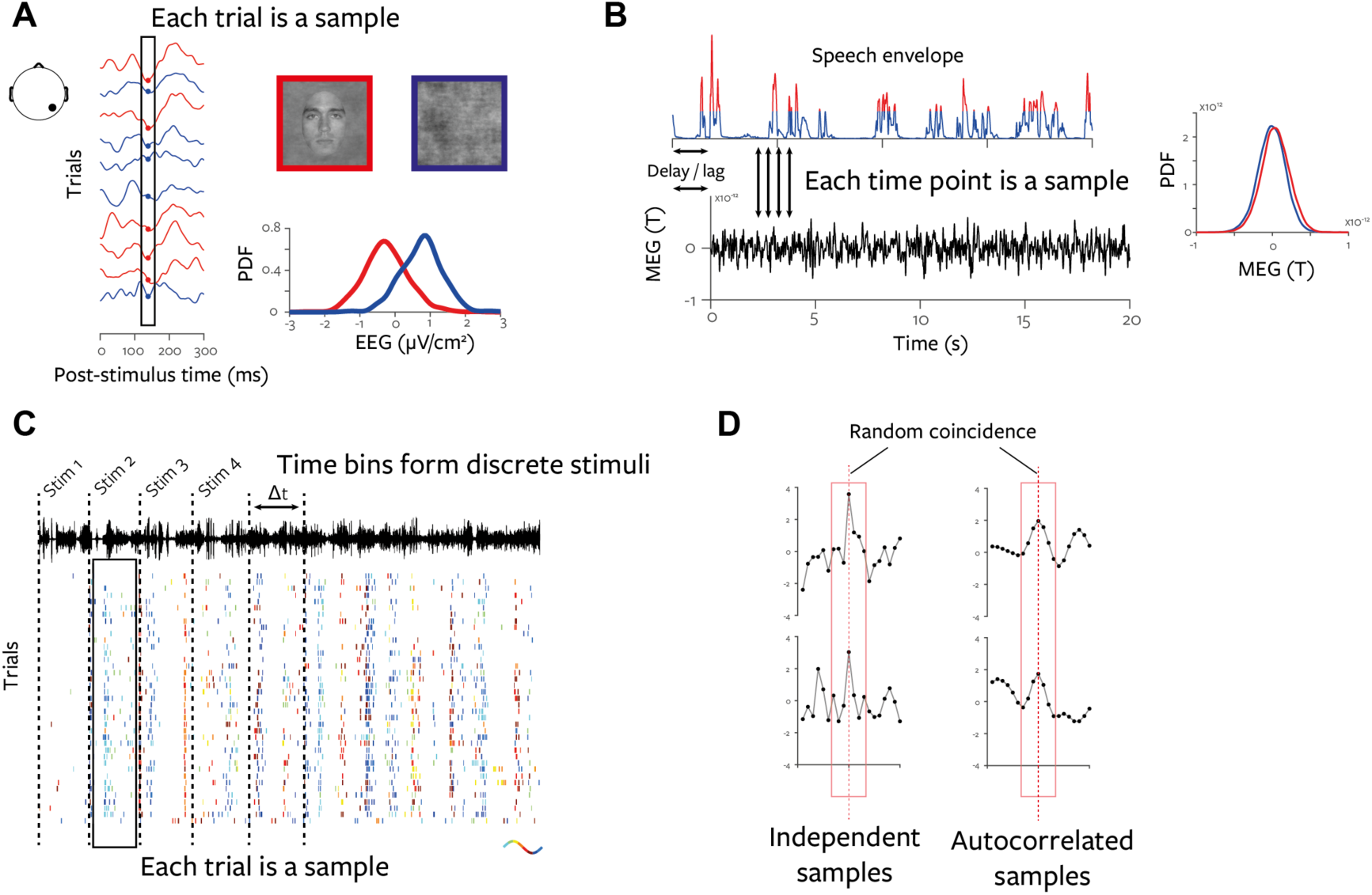
*MI calculation for different experimental designs.* Schematic illustrations show how samples used to estimate MI are obtained from different neuroimaging experimental paradigms. **A**. Event related design. Example data from a single sensor are recorded to repeated presentations (trials) of two classes of stimuli, faces (red) or noise images (blue) (Rousselet et al., 2014a). Values are extracted across presentations for a specific post-stimulus time at a specific sensor; these form the samples used for MI calculation. Kernel smoothed PDF estimates are shown for the example time point. **B.** Continuous design. Here the amplitude envelope of a speech stimulus is effectively cross-correlated with an MEG sensor signal (Gross et al., 2013). **C.** Hybrid design. A short section of a dynamic stimulus is presented many times. The stimulus is divided into bins, each of which is treated as a separate discrete stimulus. The responses over the repeated presentations (trials) of the stimulus are used as samples (Kayser et al., 2012). **D.** Two pairs of random signals with no autocorrelation (left) and with autocorrelation induced by low-pass filtering (right) are shown. The dashed red line indicates a random coincidence of high values, the red box highlights the additional relationship between the neighbouring points induced by the autocorrelation.

#### Continuous design

In a continuous design an ongoing, usually naturalistic, dynamic stimulus is presented, for example a visual movie or auditory speech. The goal of the analysis is to determine a relationship between time varying stimulus features and the M/EEG signal. The analysis is performed separately for each sensor/source and is similar to a cross-correlation. A particular delay or lag is chosen and the MI is calculated using the values of the lagged signals over time as samples (Figure 4B). Any inference requires multiple comparison correction over features, delays and sensors/sources. Since the signals usually exhibit strong autocorrelation, the permutation strategy needs to take this into account. Autocorrelation can strongly alter the distribution of the MI under the permutation null hypothesis, because if a pair of peaks in two signals happened to coincide by chance, many neighbouring points would also coincide (Figure 4D). This structure is lost if the time-domain samples are permuted without preserving the autocorrelation. We therefore suggest using a circular shifting or blockwise permutation approach when calculating MI in this sort of experimental design (Adolf et al., 2011). While there are several block-wise approaches for bootstrapping auto-correlated time series (Härdle et al., 2003; Politis and Romano, 1994), the best approach for permutation tests with neuroimaging data remains is unknown. The continuous design also imposes an implicit assumption that the neural processes under consideration are stationary for the duration of the presented stimulus.

#### Hybrid design

A third approach is a hybrid design that combined elements of both the event-related and continuous designs described above (Figure 4c). Here, a short segment of a dynamic naturalistic stimulus is presented many times. The time-course of the dynamic stimulus is split into a number of fixed-width time windows, each of which is treated as a discrete categorical stimulus. The responses obtained during that time window across the repeated presentation are used as samples for the MI calculation. This approach has frequently been applied with electrophysiology data (Strong et al., 1998) because it results in an efficient use of experimental time, and requires no prior assumptions on the specific stimulus features driving the neural response. MI calculated in this design quantifies the overall reliability of the modulation of the neural response by the stimulus without considering specific stimulus features.

As a measure of dependence MI is a function of two paired sets of samples. However, in practice sets of samples can be obtained in different ways, depending on the experimental designs just reviewed: across experimental trials, time points, or through some combination of the two. To enable meaningful interpretation of any estimated MI quantity, it is critical to properly understand how the samples were obtained via the experimental design. So, we recommend clear reporting of these design details whenever MI quantities are used.

### 2.6. Higher order information theoretic quantities

We here review some higher order information theoretic quantities and their application to brain imaging that we believe provide particularly useful and novel insights in the analyses of neuroimaging data. We describe Conditional Mutual Information, which can isolate the specific effect of a stimulus feature on neural response, while controlling the potential contribution of other correlated stimulus features; Directed Information which quantifies the time-lagged causal transfer of information between two neural responses; Interaction Information which quantifies the similarity (or synergy) of representation of the same stimulus feature between two neural responses; and Directed Feature Information which measures the time-lagged causal communication of a specific stimulus feature between two neural responses.

#### Conditional Mutual Information

Conditional Mutual Information (CMI) (Cover and Thomas, 1991) quantifies the relationship between two variables while removing any effect of a third variable. CMI between *X* and *Y*, conditioning out *Z* is usually denoted *I*(*X*;*Y* | *Z*). It is the information-theoretic analogue of partial correlation. However, while partial correlation removes only the linear effects of the third variable, CMI controls for effects of all orders and so allows for stronger conclusions to be drawn. With many types of naturalistic stimuli, extracted stimulus features are highly correlated (for example, luminance of neighboring pixels of a natural image or the acoustic features of speech). Given an analysis of each feature alone, it is difficult to determine whether a specific feature is genuinely encoded in a neural response, or whether the response is actually modulated by a different correlated stimulus feature. CMI provides a rigorous way to address this issue (Ince et al., 2012), allowing strong conclusions to be drawn about the relationship between neural responses and multiple correlated stimulus features. Figure 5 demonstrates with a simulation the use of CMI to dissociate two possible situations where a response is modulated by two correlated stimulus features.

**Figure 5.**
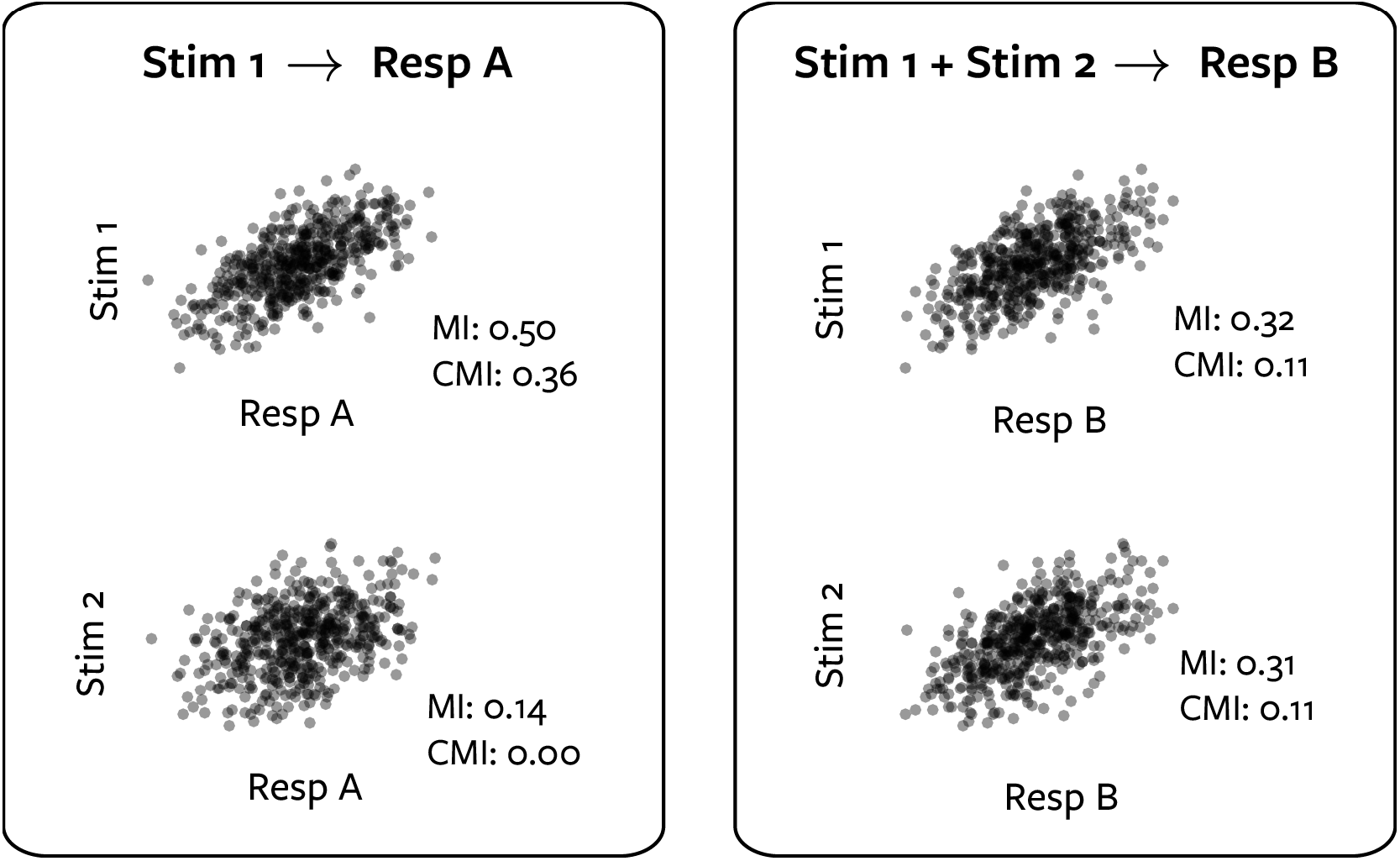
*CMI reveals genuine encoding of correlated stimulus features.* In this simulation we generated two correlated Gaussian stimulus features (covariance = 0.6), Stim 1 and Stim 2. We generated responses from two different models. Response A (left) was obtained from stimulus 1 plus Gaussian noise (top plot). In the bottom plot, MI reveals a relationship with stimulus 2 (MI = 0.14), but CMI reveals this is due only to the relationship between the features (CMI = 0). Response B (right) was obtained on each trial from the sum of stimulus 1 and stimulus 2 plus Gaussian noise. MI again reveals response dependence with both Stim 1 and Stim 2, but CMI (= 0.11) shows that now each stimulus is genuinely represented in the signal.

The ability to combine continuous and discrete variables allows for CMI to provide a novel approach to group statistics. We can construct a discrete variable representing participant identity (*P*) and use this as the conditioning variable in the definition of CMI. We can calculate the MI from the data pooled over participants (with Gaussian copula rank normalization done on a per-participant basis to account for signal differences, see Section 3.1), and also the CMI conditioned on participant identity. This CMI is the average MI effect size within each participant. These quantities are then closely related to the quantities used in the replicated G-test for independence (Sokal and Rohlf, 1981), which provides three useful inferences for group studies: First, is there a significant effect over the group (*total-G,* equivalent to CMI *I*(*S*; *R* | *P*))? This is closest to classical group inference and tells that overall the members of the group deviate from the null hypothesis (but they may not all do so and they may not deviate in the same way). Second, is there a significant difference in the effect between participants (*heterogeneity-G*, equivalent to CMI – MI, or alternatively *I*(*S*, *R*;*P*))? If so, this indicates the effect is not consistent between participants and so the data should not be pooled. This is particularly useful given that MI is an unsigned quantity – all participants could have similar levels of MI but with opposite signs of effect. It can also help to identify situations where group significance is driven by a strong effect in one or a few participants, rather than a consistent effect across the group. This is something that is not considered with most existing group statistics. Third, is the effect significant if the data is pooled (*pooled-G*, equivalent to MI)? This means overall, the data recorded across all participants deviates from the null hypothesis. This allows the identification of cases where there is a weak but consistent effect across participants, which might not suffice to produce a significant CMI value. While this approach requires further development and testing we mention it here to motivate some of the wide range of potential applications for CMI within neuroimaging.

#### Directed Information (Transfer entropy)

CMI also forms the basis for an information theoretic approach to the analysis of causal relationships between neural responses. Calculating CMI between the values of a signal *Y*, and the values of a signal *X* earlier in time, conditioning on the earlier values of *Y* itself produces a measure originally termed Directed Information (DI) (Massey, 1990) but frequently referred to as Transfer Entropy (TE) (Schreiber, 2000). DI measures the time-lagged dependence between two signals, over and above the dependence with the past of the signal itself (its self-predictability; Figure 6). This is the information theoretic analogue of Granger causality (Barnett et al., 2009; Granger, 1969) and following the arguments developed by Wiener and Granger (Granger, 1969; Wiener, 1956) can be used to infer causal relationships between brain signals, with some caveats (Bressler and Seth, 2011; Chicharro and Ledberg, 2012; Chicharro and Panzeri, 2014; Quinn et al., 2011; Wibral et al., 2014). The Gaussian copula estimate we present below (Section 3) provides a robust and computationally efficient method to estimate DI (Ince et al., 2015).

**Figure 6.**
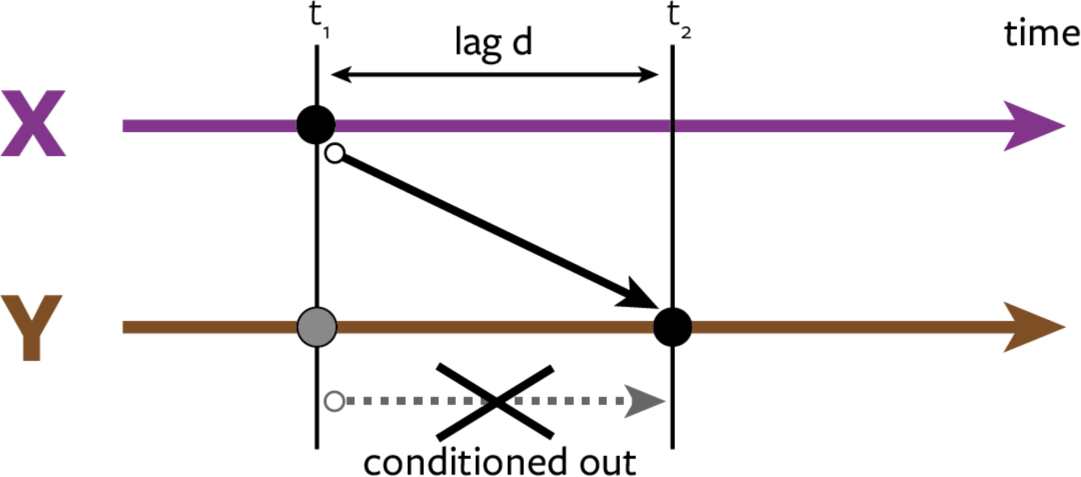
*Schematic of Directed Information (DI) calculation.* DI from *X* to *Y* is calculated as the CMI between the activity of *X* at time *t*_1_, and the activity of *Y* at a later time *t*_2_, conditioned on the activity of *Y* at *t*_1_: *I* (*X*_*t*_1__; *Y*_*t*_2__ | *Y*_*t*_1__).

#### Interaction information

It is now widely accepted that rather than operating as a number of separate functional units, the brain functions as a highly interactive distributed network. For neuroimaging studies to fully embrace this perspective we require tools to relate experimental modulations (e.g. effect of different experimental conditions) or stimulus modulations (effect of changing stimulus features) across multiple responses. For example, univariate MI analyses might reveal stimulus modulation in distinct spatial, temporal or spectral regions (Figure 7A). A natural question that arises is: Are the modulations we observe in both regions similar, or different? In other words, are the responses in both regions representing the stimuli in a similar, or a different manner?

Focusing on pairs of responses, we can schematize the stimulus MI in each response in a Venn diagram (Figure 7B). One quantity of interest is the overlap, termed the *redundancy* (Panzeri et al., 2008; Schneidman et al., 2003; Timme et al., 2013), that quantifies the MI which is shared between the two responses. Because of the additivity of MI we can obtain this quantity directly: summing the MI available in each response separately counts the overlapping region twice. We then subtract the MI available when considering both responses together, which counts the overlapping region once. The resulting value quantifies the redundancy and is equal to the negative Interaction Information (II) (McGill, 1954). Negative II corresponds to redundancy as described above, but II can also be positive, indicating *synergy* between the variables. In this case, the MI in the pair of responses when considered jointly is greater than the MI when they are considered separately. This implies that the relationship between the responses on individual trials is itself modulated by the stimulus feature considered. Redundancy is bounded above by three quantities, the MI between the stimulus and each response and the MI between the responses themselves, so it can be normalized by the minimum of these three values.

Within a neuroimaging context, high redundancy would suggest the two responses reflect the same aspects of the stimulus, and therefore likely reflect the same processing pathway or mechanisms. Alternatively, independence (zero interaction information) would suggest different processing pathways produce the observed responses. This approach can also be applied to compare different response representations or to compare responses from different experimental modalities (e.g. simultaneously recorded fMRI + EEG, Figure 7A). Similarly, interaction information can be applied in the opposite direction, considering two stimulus features (possibly from different modalities) and a single neural response, and quantifying whether they modulate the neural response in a synergistic or redundant fashion. The multivariate performance of the Gaussian copula estimate we present below (Section 3) is crucial to allow accurate estimation of the joint MI in pairs of neural responses (which themselves can be multivariate) required for computing interaction information.

**Figure 7.**
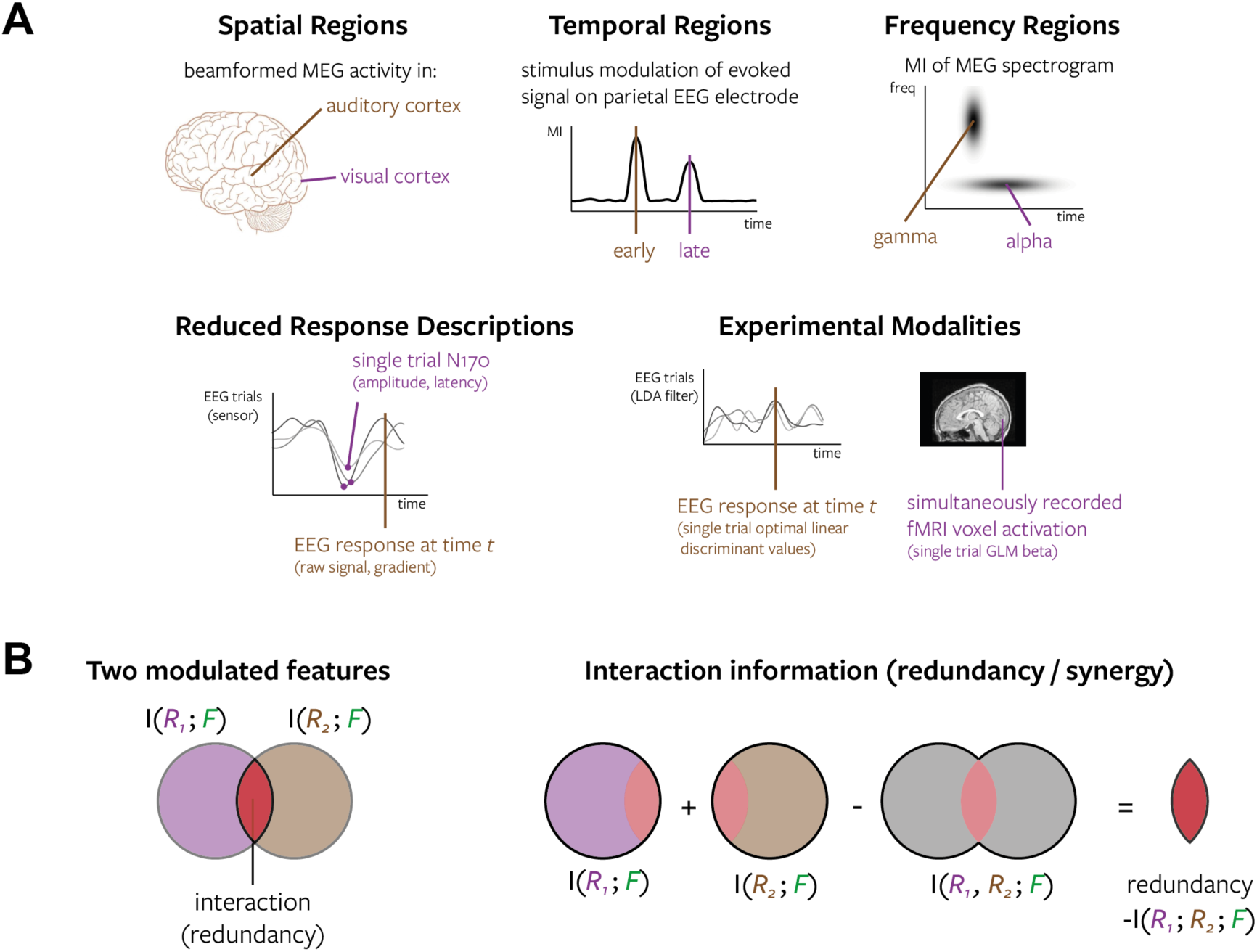
*Interaction Information: Redundancy and Synergy between neuroimaging responses*. **A**. Example situations where two different neuroimaging responses are modulated by a stimulus or task condition, and it would be of interest to relate the modulation, or information content, of the two signals. **B**. The additivity of MI allows us to quantify the redundancy (overlap) directly (see main text).

It should be noted that one issue with interaction information is that synergistic and redundant effects can cancel in the final average. While it is not clear to what degree such cancellation might occur in neuroimaging recordings this is a question that has recently received much interest (Bertschinger et al., 2014; Griffith and Koch, 2012; Harder et al., 2012; Olbrich et al., 2015; Timme et al., 2013; Williams and Beer, 2010). Further development and application of techniques to address the interplay between synergy and redundancy within a neuroimaging context is an important area for future work.

Interaction information can be applied to relate the information content of different neuroimaging responses, revealing redundant or synergistic representations. Alternative techniques to address these questions include classification images, representational similarity analysis and the temporal generalization method. Classification images (Murray, 2011) obtained from reverse correlation of different neural responses can be directly compared to quantify similarity in stimulus representation (redundancy) between areas (Smith et al., 2004). A similar approach is employed in Representational Similarity Analysis (RSA) (Kriegeskorte et al., 2008; Kriegeskorte and Kievit, 2013) which compares representational geometries between different neural responses (not the information content in the neural responses, as with classification images) by correlating dissimilarity matrices obtained from discrete category exemplar stimuli. The temporal generalization method (King and Dehaene, 2014), in which a classification algorithm is trained with neural responses from one time point and then tested on another time point, can reveal similar representations between the two time points. All of these methods are conceptually similar to redundancy, but the information theoretic approach can also reveal synergistic effects, can combine discrete and continuous stimuli or responses, can be calculated while conditioning out correlated stimulus features and can be applied to univariate responses with the full spatial or temporal precision of the considered recording modality. It also provides results with a meaningful common effect size (bits), or alternatively redundancy can be normalized to a percentage that provides an intuitive measure of the degree of overlap.

#### Directed Feature Information

We recently developed a new measure of functional connectivity called Directed Feature Information (DFI) (Ince et al., 2015). It conceptually extends DI, a measure of the amount of causal communication between two regions, to quantify the communication that is *about* a specific stimulus feature. That is, as DI measures the time-lagged relationship between the responses of two regions, DFI quantifies the amount of DI that can be attributed to variations of a given stimulus feature (e.g. the graded presence of a face in the stimulus (Ince et al., 2015)). An alternative interpretation using redundancy is that DFI quantifies the amount of redundant MI about the stimulus that is shared between *Y* and the past of *X*, over and above that which is already present in the past of *Y*. Therefore, following the Wiener-Granger principle, DFI can be used to infer the communication of the specific information about the feature considered from *X* to *Y*. DFI enables the construction of networks based on the communication of specific, task-related stimulus features rather than the networks typically constructed from the overall dependence between activity in different areas that may or may not be directly task or stimulus related. Task effects on functional connectivity can be addressed with tools such as psychophysiological interactions (PPI) analysis (Friston et al., 1997; O’Reilly et al., 2012) which reveals task induced changes in connectivity within a GLM framework. The framework of Dynamic Causal Modelling (Friston et al., 2003) can also indicate that connectivity between two brain regions is modulated by an external stimulus or condition such as attention (Penny et al., 2004). However, to our knowledge there is no other measure of functional connectivity that directly quantifies the specific content of communication as DFI does, and so it represents a transformative perspective for network-based analysis of neuroimaging data. The Gaussian copula method presented below (Section 3) is crucial to allow accurate estimation of DFI, since it requires an additional conditioning step, to calculate DI conditioned on the stimulus. The generality of the Gaussian copula estimate means this quantity can be applied to a range of situations, with discrete or continuous stimuli and potentially considering multivariate dynamic responses.

### 2.7. Relation between information theoretic quantities and other statistical approaches

Table 1 shows equivalent statistical approaches to address the same questions as the information theoretic quantities reviewed above. This illustrates how the information theoretic framework unifies a wide variety of statistical approaches with effect sizes on a common scale across many different situations.

**Table 1.** *Relation between information theoretic quantities and other statistical approaches.* Although Mutual Information (MI) is a single information theoretic quantity – a bivariate measure of dependence – here it is split into multiple rows depending on the nature of the two input variables (indicated in brackets), because different classical statistical are applicable to each of the different cases. The tri-variate information theoretic quantities below are not split by variable type – but again each of their inputs can be uni- or multi-variate and take continuous or discrete values.

| Information theoretic quantity | Other statistical approaches |
| --- | --- |
| MI (discrete; discrete) | Chi-square test of independence<br>Fishers exact test |
| MI (univariate continuous; discrete) | 2 classes: T-test, KS-test, Mann-Whitney U test<br>ANOVA |
| MI (multivariate continuous; discrete) | 2 classes: Hotelling $T^2$-test<br>Decoding (cross-validated classifier) |
| MI (univariate continuous; univariate continuous) | Pearson correlation<br>Spearman rank correlation<br>Kendall rank correlation |
| MI (multivariate continuous; univariate continuous) | Generalized Linear Model framework<br>Decoding (cross-validated regression) |
| MI (multivariate continuous; multivariate continuous) | Canonical correlation analysis<br>Distance correlation |
| Conditional Mutual Information | Partial correlation (continuous variables and linear effects only) |
| Directed Information (Transfer Entropy) | Granger causality |
| Directed Feature Information | Dynamic Causal Modeling<br>Psychophysiological Interactions |
| Interaction Information | Representational Similarity Analysis (redundancy only)<br>Cross-classification decoding (redundancy only)<br>Mediation analysis |

## 3. A novel method for MI estimation using a Gaussian copula

### 3.1. Estimating MI between two continuous variables with a Gaussian copula

We present here a new estimator of MI that uses the concept of a statistical copula to provide the advantages of Gaussian parametric estimation (Section 2.4) for variables with any marginal distributions. A copula (Nelsen, 2007) is a statistical structure that expresses the relationship between two random variables, independently of their marginal distributions. Sklar’s theorem (Sklar, 1959) states that any multivariate distribution can be expressed as the combination of univariate marginal distributions and an appropriate copula – the copula links the individual variables and represents the statistical relationships between them. Copula based analyses have been widely applied in quantitative finance (Genest et al., 2009a) and have recently been applied to estimate Granger causality in neuroimaging data (Hu and Liang, 2014).

Formally, Sklar’s theorem states that every multivariate cumulative distribution function *F*(*x*_1_,…,*x*_*k*_) = *P*(*X*_1_ ≤ *x*_1_,…,*X*_*k*_ ≤ *x*_*k*_) can be expressed in terms of its marginal cumulative distribution functions *F*_*i*_(*x*) = *P*(*X*_*i*_ ≤ *x*) and a copula function *C*:[0,1]^*k*^ → [0,1]:

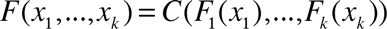

If the marginals *F_i_* are continuous then the copula C is unique.

Figures 8A and 8B illustrate the copula concept with simulated Gaussian data for uncorrelated and correlated variables respectively. Left hand scatter plots show 1000 simulated data points. Right hand scatter plots show the empirical copula of this simulated data: the empirical cumulative distribution function (CDF) value of each variable evaluated at each data point. The empirical CDF is calculated by ranking each sample (separately for each variable) and then rescaling the integer rank values to the range (0,1). Thus, the resulting bi-variate distribution (copula) is a probability density over the unit square. For independent variables (Fig. 8A) the copula is uniform, while for correlated variables (Fig. 8B) the copula has non-uniform density.

**Figure 8.**
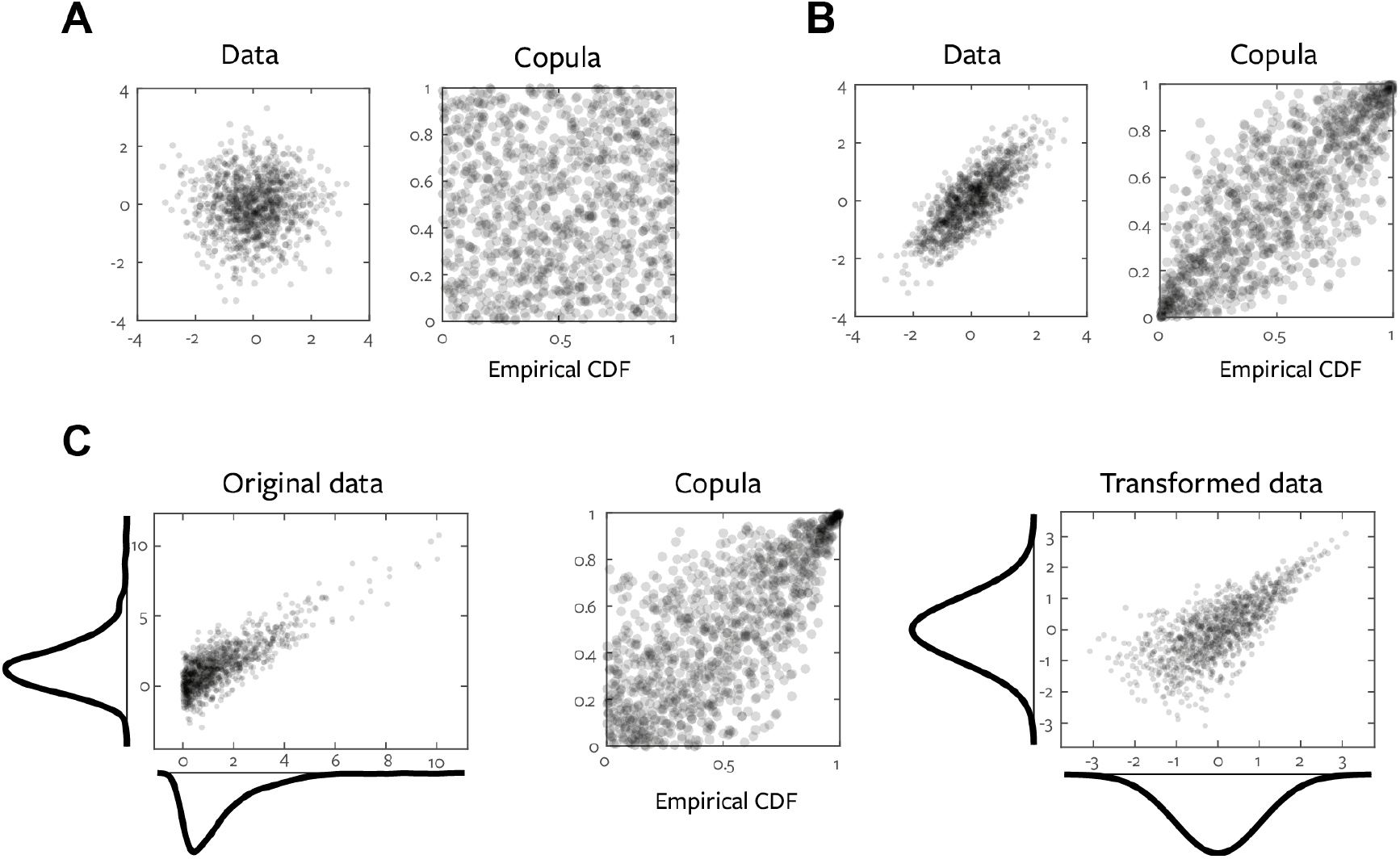
*Examples of Gaussian copulas and a copula-preserving Gaussian marginal transformation.* **A**. Scatter plots of simulated data from two independent standard normal variables, and their copula. **B**. Scatter plots of simulated data from two correlated standard normal variables (r=0.8), and their copula. **C.** Scatter plot and marginal densities of data simulated from the model x=exp(1.5); y=x+N(0,1) (left), the empirical copula (center) and the transformed data with Gaussian marginal but the same empirical copula as the original data (right).

This is useful because the copula linking two variables is directly related to the MI between them. The entropy of the joint distribution over two variables, *X* and *Y*, is equal to the marginal entropies plus the entropy of the copula:

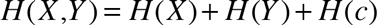

where *H* (*c*) is the entropy of the copula density, *c*, which links *X* and *Y*.

Plugging this into the third form of Eq. (1), the marginal entropies cancel revealing that the MI between X and Y is equal to the negative entropy of their copula (Calsaverini and Vicente, 2009; Kumar, 2012; Ma and Sun, 2011; Zeng and Durrani, 2011); thus the copula fully encapsulates the relationship between the two variables. A corollary of this is that MI does not depend on the marginal distributions of the individual variables.

We can therefore estimate MI by applying continuous entropy estimates to the empirical copula density (Ma and Sun, 2011); however these can still be computationally and data intensive. Instead we exploit the corollary mentioned above; since the copula entropy, and hence the MI, does not depend on the marginal distributions of the original variables, we can transform the marginals in any way we see fit. As long as we preserve the empirical copula linking the variables, the statistical relationship that is quantified by MI will be unchanged. We therefore transform the marginals to be standard Gaussian variables, to which we can apply the efficient parametric MI estimate described in the previous section.

Figure 8C illustrates this transformation; the variable plotted on the x-axis is drawn from an exponential distribution (rate=1.5); the variable on the y-axis is that value added to a standard normal. The left hand scatter plot of 1000 samples from this model shows the non-Gaussian marginal distributions of these data and the copula plot (center) illustrates the dependence between the variables. For each sample, the transformed value of each variable is obtained as the inverse standard normal CDF evaluated at the empirical CDF value of that sample. By the probability integral transform, the empirical CDF values of the data sample are uniformly distributed and so this transformation produces a data set that has perfect standard normal marginals. This transformed data set preserves the same empirical copula as the original data (right); in other words, the rank-relationships between the variables are preserved. In practice, the empirical CDF is not computed explicitly; instead the rank of each sample is obtained and normalized by *N+1*, where N is the number of samples. This results in a uniform distributed sample taking values in the range 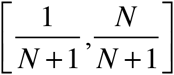 at which the inverse standard normal CDF is directly evaluated. Transformations of this type are sometimes referred to as Inverse Normal Transformations and have been used in fields such as genetics (Beasley et al., 2009). We can then calculate MI between the transformed variables using the parametric Gaussian model (Eqs 1,3).

The parametric MI estimation implicitly imposes the assumption of a Gaussian copula linking the two variables. If this assumption is violated the resulting MI value may not be accurate. However, since the Gaussian distribution has the maximum entropy for a given mean and covariance (Cover and Thomas, 1991) the Gaussian copula must also have the maximum entropy of possible copula models preserving second order statistics. Otherwise the distribution of the same two Gaussian variables linked by this higher entropy copula would itself have a higher entropy, contradicting the proven maximum entropy property of the Gaussian. Since MI is the negative copula entropy other choices of parametric copula models (or direct estimation) could give higher, but not lower MI estimates: the Gaussian copula estimate is therefore a lower bound to the true MI value (Calsaverini and Vicente, 2009). This lower bound property is crucial for an estimator that is to be used for statistical testing, since it ensures that erroneous high values cannot occur due to mismatched assumptions between the statistic and the data; the measured value is always lower than the true value. In the multivariate case, the same Gaussian marginal transformation is applied independently to each constituent variable. This preserves the rank relationships both within and between the two multivariate variables considered for the MI calculation (X and Y in Eq. 1). Zeng and Durrani (2011), propose to estimate MI between univariate variables via a Gaussian copula, the entropy of which is estimated via Kendall’s tau. The advantage of the approach presented here is that it can be applied to multidimensional variables (see Section 4) and to estimate MI between discrete and continuous variables (Section 3.2).

In summary, by transforming each univariate marginal to be a standard normal and applying a Gaussian parametric MI estimate, we obtain a lower-bound estimate of the MI. We call this estimator Gaussian-Copula Mutual Information (GCMI). As the value of this estimate depends only on the empirical CDF of the data, it is in effect a rank statistic and so robust to outliers. Although the estimate derives from a parametric assumption on the copula linking the two variables, there is no assumption made on the marginal distributions and it can therefore be applied to any continuous valued data. Within neuroimaging, we propose use of this estimator as a test statistic for a permutation based hypothesis test with approaches that include correction for the problem of multiple comparisons (see Section 4). We therefore focus primarily on this application in this paper (Sections 4.1-4.3). However, we address in more detail the bias of the estimator and the implications of the lower bound property in Section 4.4.

### 3.2. Estimating MI between a discrete and a continuous variable using a copula based approach

In many cases, the statistical inference of interest concerns a continuous valued neuroimaging signal recorded in response to a number of different discrete stimuli or under different experimental conditions. Here we extend the copula-based MI estimate introduced above to this problem – estimating MI between a (univariate) discrete and a (potentially multivariate) continuous variable. Despite the wide applicability of such a measure, the practical issue of computing MI between discrete and continuous variables has so far received little attention (Lefakis and Fleuret, 2014; Magri et al., 2009; Ross, 2014). One approach would be to discretize the continuous variable as described in Section 2.3 and use standard binned methods. However as discussed previously this suffers from the curse of dimensionality when considering multivariate spaces. Instead we develop an approach based on the Gaussian copula estimator we described above.

Given the lack of explicit parametric distributions over mixed continuous and discrete spaces it is convenient to start from the second form of Eq. (1). With *X* as the (possibly multivariate) continuous neuroimaging response and *Y* as the discrete stimulus, we have from the definition of conditional entropy 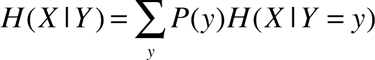 and so:

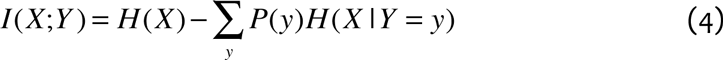

It is straightforward to apply Gaussian parametric entropy estimation to the conditional entropy terms, using the samples available for each *y* value. However, when considering the unconditional entropy term *H*(*X*) there are two possible approaches. One option is to form the mixture model from the class conditional parametric fits and numerically integrate this to obtain the entropy (since there is no closed form expression for the entropy of a Gaussian mixture). A second option is to fit a separate parametric model of the same form to the full data set and use that to estimate the unconditional entropy. Figure 9 illustrates these issues with an example of data generated under a Gaussian model with different means for each of two discrete stimulus conditions (Figure 9A). Figure 9B illustrates the two different models that can be used to estimate the unconditional entropy term, the actual mixture density (solid line) or a Gaussian fit (dotted line). Whichever strategy is used, MI is obtained as the difference between the entropy of the chosen unconditional model, and the average of the entropies of the class conditional distributions (Figure 9C). It is not clear if either of these is more appropriate than the other – each preserves a different interpretation of MI. The first leads to an MI estimate more consistent with the Kullback-Leibler divergence expression of MI; it is the property of a single distribution (the fitted parametric mixture distribution) and measures the deviation of that distribution from a surrogate independent model. The second leads to an MI estimate more consistent with a statistical testing viewpoint, comparing an unconditional Gaussian fit to class-conditional Gaussian fits (c.f. ANOVA).

**Figure 9.**
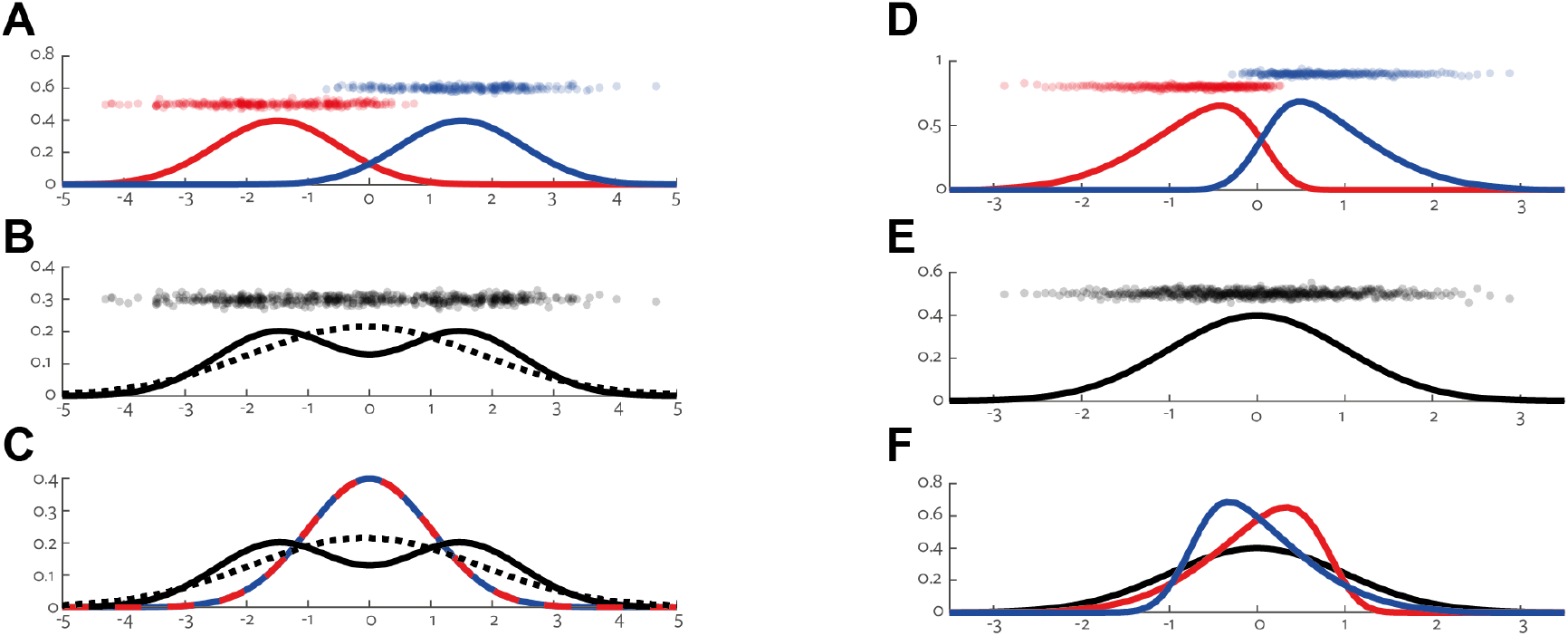
*Illustration of continuous-discrete MI calculation*. Left hand panels show data generated from a Gaussian mixture model with two classes. **A**. Sampled data points and PDF for each class. **B**. Sampled data points, unconditional PDF of the data (solid line), and maximum likelihood Gaussian fit (dotted line). **C**. True unconditional PDF (solid line), unconditional Gaussian fit (dotted line) and de-meaned class-conditional PDF’s (red/blue). Right hand panels show the same data after copula Gaussian transformation. **D**. Transformed sampled data points and PDF (kernel density estimate) for each class. **E**. Transformed sampled data points and unconditional probability density (solid line). **F**. Unconditional PDF (black) and class-conditional PDF’s (red/blue).

Fortunately, the use of the copula transform removes this dichotomy. Motivated by the previous section we apply the same copula transform to the unconditional data to obtain a surrogate data set with a standard normal distribution (Figure 9E), while preserving the rank-class relationships. Figure 9D shows the resulting class conditional distributions. Again, MI is calculated as the difference between the entropy of the unconditional entropy and the average entropy of the class conditional distributions (Figure 9F; Eq. (4)). Now, by design the true unconditional distribution is Gaussian, so the unconditional Gaussian fit and the class-conditional mixture are equivalent.

It is clear that after the transformation, the class-conditional distributions are no longer Gaussian (Figure 9D), so our Gaussian conditional entropy estimate will be an approximation. However, again the maximum entropy property works in our favor: each class conditional Gaussian entropy estimate will necessarily be greater than or equal to the true entropy of the class. Since the conditional entropy terms are subtracted in Eq. (4) this ensures the estimated MI is again a lower bound on the true MI. As in the continuous case, this method can be applied whatever the original distribution of the data; it does not require Gaussian classes as in the example shown here. The key feature is that the copula transform preserves the rank-class relationships, and results in a data set to which the parametric Gaussian entropy estimates can be applied. Note that while the unconditional entropy *H(X)* itself is not invariant to the copula normalization transform, as in the continuous case, the mutual information, as a difference of entropies, is invariant to marginal transformation.

### 3.3. Estimating MI in spectral data: phase and power

In many applications, analyzing M/EEG signals in the frequency domain is of particular interest because of the potential for understanding brain oscillations, which are increasingly thought to underlie many important cognitive processes (Schnitzler and Gross, 2005; Singer, 2013; Thut et al., 2012; Wang, 2010). The methodological issues surrounding how best to extract a frequency-based representation from M/EEG data, and perform statistical analysis on such data to determine the presence and nature of stimulus modulations have therefore received much attention (Gross, 2014). In this section, we emphasize how our new Gaussian copula MI estimate can be applied to spectral data. In fact, the approach described here can be applied to any vector quantity (e.g. magnetic field vectors) to separate stimulus modulations of amplitude and direction.

The most commonly employed frequency or time-frequency decompositions result in a complex valued spectral signal in each frequency band (Gross, 2014). The real and imaginary parts of this complex signal can be treated as a 2D response variable within the Gaussian-copula MI framework, where each is transformed separately prior to parametric MI estimation. This can be applied with either the continuous-discrete or continuous-continuous MI as described previously, to allow quantification of categorical experimental differences, as well as encoding of continuous valued stimulus features.

There is often additional interest in characterizing more specifically which aspects of the oscillatory activity – amplitude (the size of the oscillations) or phase (the temporal alignment of the oscillations) – are modulated by experimental conditions or activity in other brain signals (frequency bands or regions). Binned MI measured have already been applied to these sorts of problems (Belitski et al., 2010; Gross et al., 2013; Kayser et al., 2009; Schyns et al., 2011; Szymanski et al., 2011), and a number of other techniques have also been developed (Kempter et al., 2012; Lachaux et al., 1999; Voytek et al., 2013). In such analyses, it is unclear how best to model or bin the data, because phase, the angle of the complex spectral signal, is a circular variable which ‘wraps around’ and has no clear ranking or extremal values (Berens, 2009; Lee, 2010). One example is whether the arbitrary cut-off in the numerical representation used should be taken as a bin edge (usually angles are returned in the range [−π, π] but this is implementation dependent).

With the Gaussian copula MI estimator, we suggest extracting phase and amplitude of a complex spectral signal as follows (Figure 10). We obtain amplitude in the normal way as the absolute value of the complex number. This is then transformed as a 1D variable for use with a Gaussian-copula MI estimate (Figure 10; center amplitude plots). Often the square of this amplitude is used and referred to as power, but we note that within the copula framework MI is a rank statistic and since amplitude is always positive and the square operation is monotonic, the choice of power or amplitude does not affect the MI value. To isolate phase, we normalize the complex number by its amplitude, resulting in a 2D variable where all points lie on the unit circle. We then apply the 2D Gaussian copula MI estimate on this 2D variable; transforming each dimension independently (Figure 10, right hand phase plots). Maintaining a 2D representation for phase avoids the technical issues surrounding circular variables, particularly modeling joint distributions of circular and linear variables. The Data Processing Inequality ensures that since only information about phase goes into the calculation (all amplitude variation is removed), whatever processing we apply (here copula transformation) cannot add information and so our estimate of MI carried by phase is valid. Figure 10 illustrates the approach with two simulated systems, one in which only phase is modulated by a discrete experimental condition (Figure 10A), and one in which only power is modulated (Figure 10B). The bar graphs of MI in the different signal representations (Figure 10; far right) demonstrate that this approach can correctly dissociate the two types of modulation.

**Figure 10.**
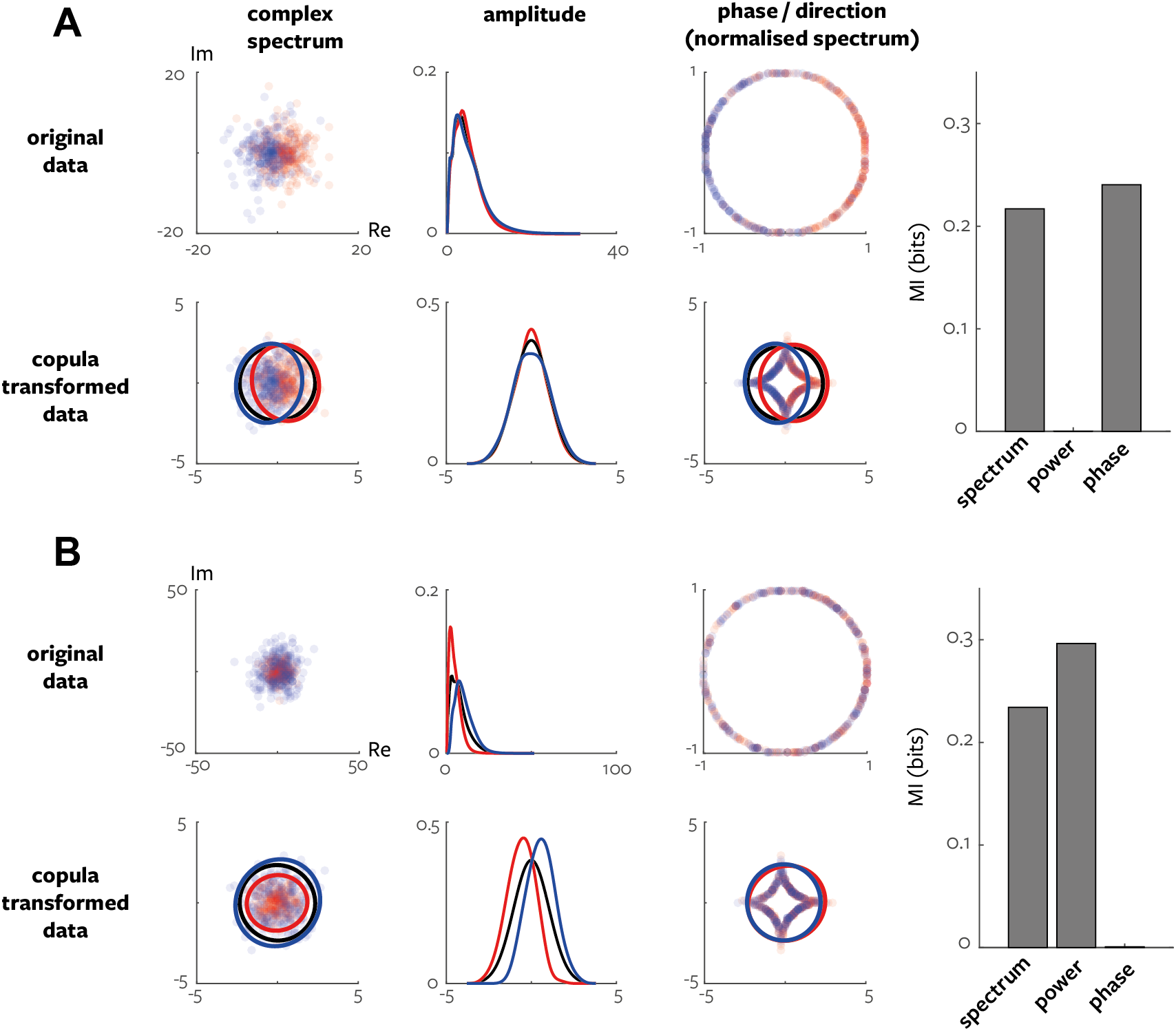
*Gaussian-copula MI applied to complex spectral data.* Spectral data were generated from two two-class models. **A**. Phase was sampled from von Mises distribution with class specific mean and amplitude was sampled from chi-square distribution (common across classes). **B**. Phase was drawn from von Mises distribution (common across classes) and amplitude was sampled from chi-square distribution with class-specific degrees of freedom. **A,B**. Left plots show generated complex data (top) and with marginal copula transformation (bottom). Solid lines show p=0.01 contours of multivariate Gaussian pdf. Centre plots show amplitude (top) and copula transformed amplitude (bottom). Right plots show amplitude-normalized spectrum (top) and copula transformed normalized spectrum (bottom). Far right bar graphs show the MI value in the different data representations.

We emphasize that our approach is equally applicable to the single-trial outputs of any frequency or time-frequency decomposition including Empirical Mode Decomposition, Hilbert-Huang transform and matching pursuit methods (Gross, 2014). It can also be applied to other vector quantities to determine the relative effects on amplitude and direction, for example planar magnetic field gradients (see Section 4.2).

## 4. Results

Our primary intention is to present our new Gaussian-Copula Mutual Information (GCMI) estimator as the effect size for a practical statistical test for neuroimaging, that can be considered as a drop-in replacement for a number of different established statistical measures (Table 1). In this section we therefore first validate the performance of the new estimator when employed as a statistical test and provide some example applications (Sections 4.1-4.3). The data used for the simulation and examples in this section are available in (Ince et al., 2016). We then demonstrate the bias and mean-square error of the GCMI estimator compared to other mutual information estimators on simulated systems as well as the example data sets (Section 4.4).

### 4.1. Discrete experimental condition with continuous EEG response: Face detection

We performed a number of analyses in order to evaluate the performance of the Gaussian copula continuous-discrete MI estimator (Section 3.2) as a statistical test for neuroimaging applications. We consider EEG data collected from a single subject within an event-related design (Section 2.3), with presentation of two classes of images: faces and noise textures. Data were band-pass filtered between 1-30Hz and the Current Source Density transformation was applied (Rousselet et al., 2014a). We calculated MI independently for each time point and sensor using samples collected from the repeated presentations (Figure 4A). A common approach with neuroimaging studies is to apply a permutation test together with the method of maximum statistics in order to correct for multiple comparisons (Holmes et al., 1996; Nichols and Holmes, 2002). We calculated the two-sample Kolmogorov-Smirnov (KS) test statistic (Massey, 1951) for each time point and sensor from all the available 1000 trials and used this as the ground-truth to evaluate tests performed with smaller numbers of trials (Figure 11A, top left, color plot). We performed the calculation over all sensors and time points 1000 times, randomly permuting the stimulus class labels each time. We took the maximum value over sensors and time points for each of these permutations, and used the 99^th^ percentile of these maximum values as the threshold for significance (Figure 11A, top left, black and white plot). This procedure corrects for multiple comparisons and provides a Family-Wise Error Rate (FWER) of 0.01. We then repeatedly sub-sampled smaller sets of trials from the full data set and repeated the mass-univariate analysis for various statistics, together with the permutation approach. Figure 11A, middle and bottom show the results for a sub-sampled set of 100 trials. The significant sensors and time points were then compared to the full data KS-test ground truth to evaluate the performance of different statistics.

We considered four types of statistic, the new Gaussian copula MI, binned MI (with 2,4 or 8 bins), the t-test (unequal variances) and the KS-test. Figure 11B shows the time-courses obtained from a single sensor for each of these statistics. First we considered the statistical performance of the null-hypothesis test based on each of these statistics over the full space of time points and sensors, as a function of the number of trials available. For each sample size (25, 50, 100, 200, 500), we randomly selected that number of trials from the full data set, calculated all the statistics including 200 class-shuffled permutations, and determined the final multiple-comparison corrected inference for each statistic. We repeated this procedure 50 times for each sample size. For each statistic and sample size we then compared the result of the inference to the ground truth, considering sensitivity (Figure 11C, top left), specificity (Figure 11C, middle left) and MI with ground-truth (Figure 11C, bottom left). Sensitivity, or true positive rate, measures the proportion of significant ground-truth responses that are correctly detected with each test statistic. Sensitivity increases with number of samples for all statistics, the t-test and copula MI have the highest sensitivity over the full range of trials considered. Specificity, or true negative rate, measures the proportion of non-significant ground-truth responses that are correctly detected as non-significant by each test. Specificity is high for all statistics., due to the strong control on FWER provided by the permutation approach. Binned MI methods have the highest specificity, the t-test has the lowest, with the copula MI taking intermediate values. Finally, as an overall measure of the performance of the test statistics we consider the MI in the contingency table (or confusion matrix) for each test (Quian Quiroga and Panzeri, 2009); i.e. for each repetition, the discrete MI between the binary ground-truth significance and the significance for that repetition, with time points and sensors providing the samples for the MI calculation. Similar in spirit to Matthew’s Correlation Coefficient (Baldi et al., 2000; Matthews, 1975), this provides an overall accuracy measure, which incorporates possible asymmetries resulting from unbalanced classes. With this measure, all test statistics perform better with more samples; the t-test and copula MI stand out as more accurate than the other tests, with the t-test the most accurate.

Next, we fixed the number of trials at 100, and performed a similar analysis to evaluate the robustness of the different statistics to the presence of outliers. On each of 50 repetitions, a certain percentage of trials were chosen (0-50%) and for those trials the EEG signal at each sensor and time point was replaced with a random variable drawn from a Gaussian distribution with standard deviation (s.d.) 5 times greater than the actual s.d. of that response. We again computed sensitivity, specificity and MI in the confusion matrix against the same undistorted 1000 trial KS-test ground truth (Figure 11C, right). The sensitivity and MI plots illustrate the robustness of the copula MI statistic compared to the t-test. While the t-test has slightly higher sensitivity for the original data, 5% of trials corrupted by outliers is already enough to reduce the sensitivity of the t-test to half that of the copula MI. The KS test shows higher robustness than the GCMI based test when there is a lot of noise (>10% of trials corrupted by noise). However, at low noise levels it is less sensitive (GCMI detects ~60% more true positives in the original data with no corrupt trials). A similar pattern is seen when comparing the binned MI methods to GCMI – reduced sensitivity at low noise levels, but increased robustness at high levels of noise.

**Figure 11.**
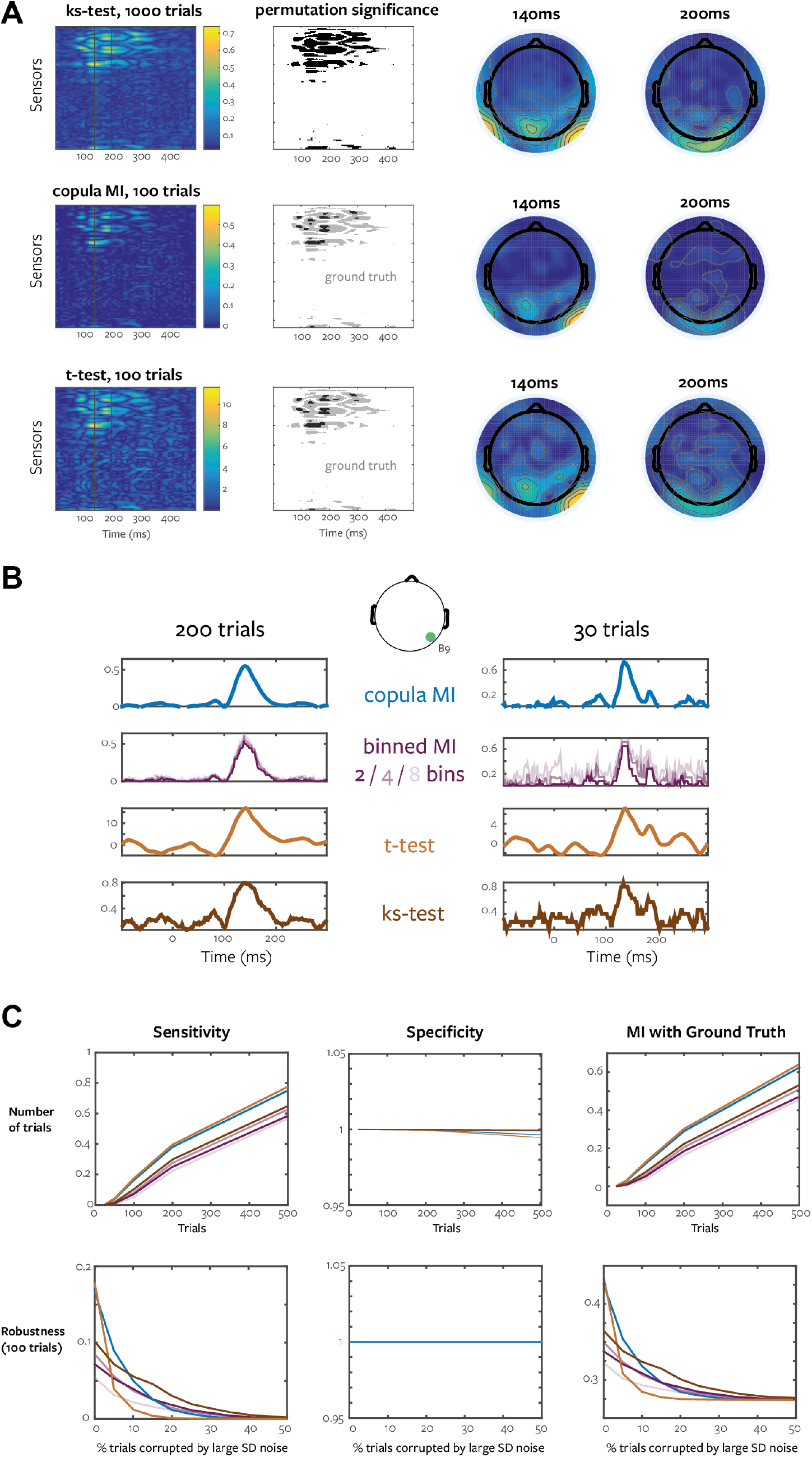
*Performance of Gaussian copula MI estimator as a statistical test for EEG data with discrete stimuli in event related design.* **A.** Statistics are calculated for each sensor and time point (left colored image plots) and significance determined with permutation testing and the method of maximum statistics (black and white image plot). Topologies are shown for two indicated time points. The KS-test with all 1000 trials is used as the ground-truth (top); copula MI with 100 trials (middle) and t-test with the same 100 trials (bottom) are shown. The results of permutation significance for these statistics are shown (black) overlaid on the ground-truth significance (grey). **B.** Example time-courses of various effect sizes calculated with 200 (left) or 30 (trials). **C.** Results of numerical investigation of the performance of various statistics with permutation testing, as a function of the amount of data available (left column) and as a function of the amount of noise added to the data (right column).

In summary, when performing mass-univariate analyses with permutation testing and maximum statistics, the copula MI provides similar sensitivity to the more commonly employed t-test, but is considerably more robust to the presence of outliers. It also performs better than binned MI estimates, and can be applied to multi-variate responses.

### 4.2. Continuously varying experimental condition with continuous MEG response: listening to speech

To evaluate the performance of the Gaussian copula continuous-continuous MI estimator (Section 3.1) we consider MEG data collected from a single subject within a continuous design with an auditory speech stimulus (Gross et al., 2013; Park et al., 2015). For simplicity we focus here on a sensor-level time-domain analysis. Since previous work has shown speech entrainment mostly at lower frequencies, we extracted the wideband amplitude envelope of the speech stimulus (Chandrasekaran et al., 2009; Gross et al., 2013) and then low-pass filtered with a 12 Hz cutoff (3^rd^ order non-causal Butterworth). The MEG signal was obtained from a 248-magnetometer whole-head MEG system (MAGNES 3600 WH, 4-D Neuroimaging). We band-pass filtered in the range 2-12 Hz (3^rd^ order non-causal Butterworth), downsampled to 50Hz and then computed the planar gradient tangential to the head at each magnetometer (Bastiaansen and Knösche, 2000). This potentially simplifies interpretation of sensor-level data because it typically results in maximal signal directly above the corresponding source (Hämäläinen et al., 1993). We analyse 450s of recording (22,500 samples at 50Hz) during which the subject listened to a spoken story. Similar to a cross-correlation of two signals we calculate the relationship between the speech envelope and the MEG signal over a range of delays (0-350ms).

**Figure 12.**
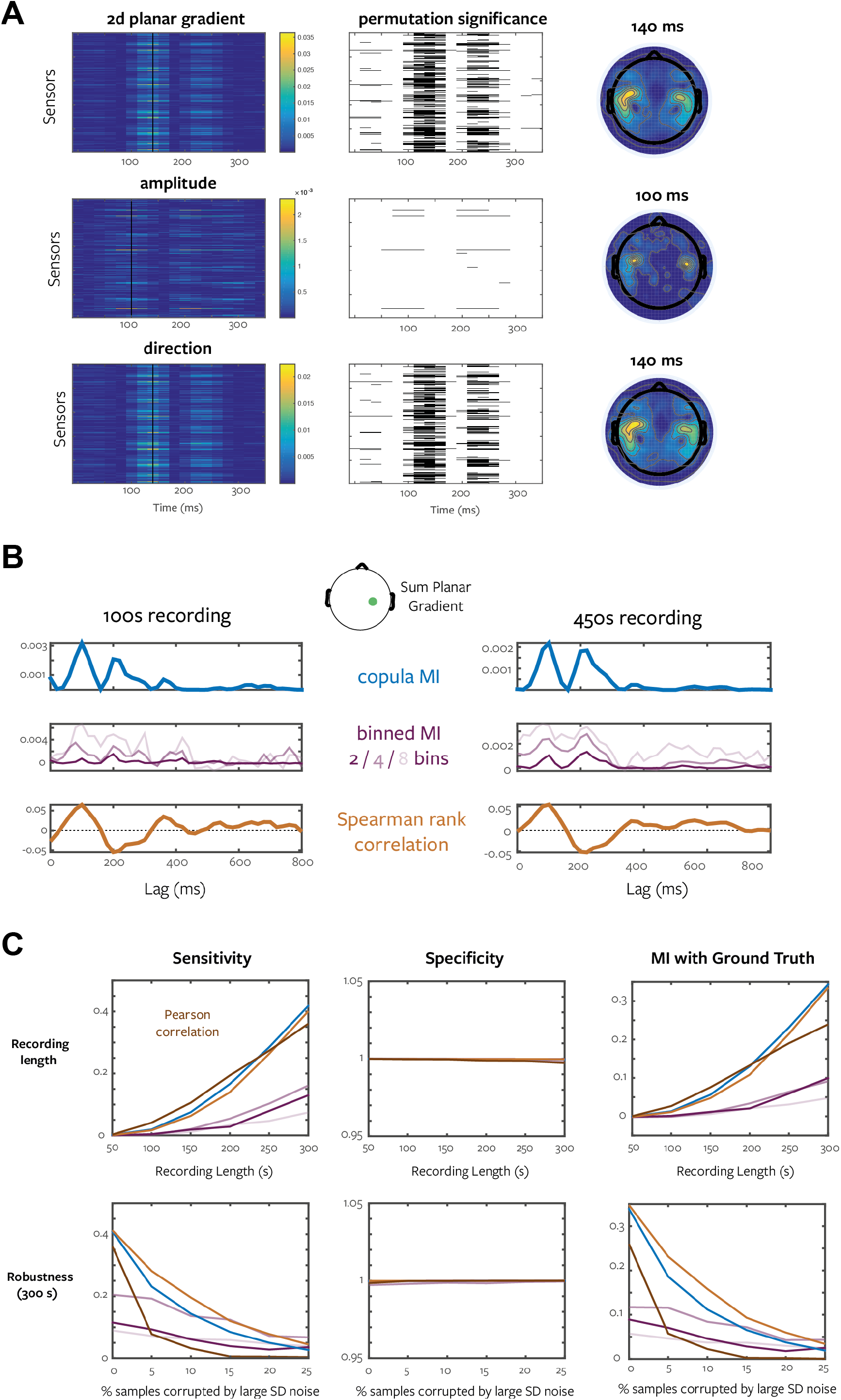
*Performance of Gaussian copula MI estimator as a statistical test for MEG data with continuous stimuli in a continuous design.* **A.** Gaussian copula MI is calculated between the speech envelope and the full 2D planar gradient response (top), the amplitude (middle) or the direction (bottom) of the planar gradient vector, for each sensor and time point (left colored image plots). Significance is determined with block permutation testing and the method of maximum statistics (black and white image plots). Topologies are shown for the indicated time points. **B.** Example cross-correlation style delay plots of various effect sizes calculated with 100s or 450s of continuous stimulation. **C.** Results of numerical investigation of the performance of various statistics with block permutation testing, as a function of the amount of data available (left column) and as a function of the amount of noise added to the data (right column).

The resulting planar gradient signal consists of a 2d magnetic field vector (tangential to the head) at the position of each magnetometer. Typically, the amplitude of this 2d vector is used as the response signal of interest (Oostenveld et al., 2011). However, using the multivariate MI estimate we can quantify the modulation of the full 2d signal, as well as breaking down the stimulus effects on amplitude and direction separately (Figure 12A), as described for spectral data in Section 3.3. The top row of Figure 12A shows the MI between the speech envelope and the 2d planar gradient for each channel and speech-MEG delay lag. The black and white image plot shows the multiple comparison corrected permutation significance (p=0.01, 200 permutations, 10s block permutation scheme to preserve signal autocorrelation). The middle row shows the same for the 1d Pythagorean amplitude (Fieldtrip default), and the bottom row shows the MI in the planar gradient direction only, with the amplitude effects normalised out as described for phase in Section 3.3. This shows that, while there is a focal and statistically significant modulation of the planar gradient amplitude over the auditory cortices, in fact the direction of the planar gradient is modulated much more strongly by the speech envelope over a much wider area, with MI values an order of magnitude higher. In addition, the timing of the peak effect is different (earlier for the amplitude). This suggests that focusing on the amplitude of time-varying magnetic field vectors could result in reduced sensitivity, and provides an example of the potential advantage of using multivariate statistics that allow separate treatment of the direction and amplitude of vector values.

However, to investigate the properties of the copula MI estimator when employed within a permutation based null-hypothesis testing framework we use the 1d amplitude signal, as there are not so many well-established statistical methods to compare for evaluating the relationships in the continuous multivariate case. Figure 12B shows the delay time courses for the sensor with the strongest amplitude modulation. To determine the performance of the copula MI as a statistical test we proceed as described in the previous section. First, we obtained ground-truth significance by applying Spearman’s rank correlation between the speech envelope and the lagged MEG planar gradient amplitude using the full 450s of available data (1000 permutations, 10s block permutation scheme, maximum statistics corrected over all sensors and 18 delays considered). Then we subsampled (using the 10s block scheme) reduced amounts of data (50-300s, repeated 30 times each). For each subsampled repetition we computed a range of statistics (copula MI, binned MI, Pearson correlation, Spearman correlation) and determined the significance of each with block permutation and maximum statistics (p=0.05, 100 permutations, 10s blocks) (Figure 12C, left column). Sensitivity, specificity and MI with ground-truth show that the copula method performs similarly to rank correlation (slightly lower sensitivity, slightly higher MI with ground-truth).

To investigate the robustness of the statistics we fixed the amount of data at 300s and investigated the effect of corrupting a fixed percentage of trials with Gaussian noise with standard deviation 5 times larger than that of the MEG signal (Figure 12C, right column). Here Spearman rank correlation performs best (measured both with sensitivity and MI), but the copula MI is close. Pearson correlation is very sensitive the addition of outliers with sensitivity reduced to ~30% of that of the copula MI with only 5% corruption. The use of Spearman’s correlation to define the ground-truth, combined with the low number of significant responses may result in a bias towards the particular property of that measure.

In summary, when performing mass-univariate analyses with permutation testing and maximum statistics, the continuous copula MI provides similar performance as Spearman rank correlation. However, as an MI estimate it benefits from the useful properties of the MI effect size (e.g. additivity; see below), and we have shown that the ability to consider multivariate responses and separately quantify modulations of vector direction and amplitude have the potential to provide more detailed interpretations of MEG data.

### 4.3. Pairwise temporal interactions reveal modulation of gradient in EEG

To provide an example application of interaction information (Section 2.6; Figure 7) we consider temporal interactions within an event-related EEG experiment with a continuous valued stimulus feature. The task was face detection, with stimuli as in Figure 4A, but here the images were sampled with Bubbles: randomly positioned Gaussian apertures which selectively reveal different parts of the image on different trials (Gosselin and Schyns, 2001). Since it has been shown that in this paradigm it is primarily the visibility of the eye region which modulates the recorded EEG (Rousselet et al., 2014a), we focus here on the visibility of the left eye region (a continuous scalar value for each trial obtained by summing the bubble masks within the eye region) and its effects on a contralateral right occipital-temporal electrode. We consider 1092 trials from a single observer during which a bubbled face image was presented (Rousselet et al., 2014b). EEG data were band-pass filtered (1-30Hz) and the Current Source Density transformation was applied (Rousselet et al., 2014a).

Figure 13D shows how the modulation of the evoked time-course by the stimulus feature (eye visibility), and how this can be quantified by calculating Spearman’s rank correlation or copula MI independently at each post-stimulus time point (Figure 4A). However, with MI we can calculate the Interaction Information (II) between pairs of time points (Figure 13A). This allows us to investigate the relationship between the modulation of the evoked signal at different times: positive II indicates a synergistic relationship between the responses; negative II means they are redundant. Here, the MI curve has 3 peaks; the temporal interaction matrix reveals that the second and third peaks are mutually redundant, but the first peak appears to carry independent MI. Interestingly, there are striking patches of local synergy (indicated with dashed lines), equivalent in magnitude to the largest MI values over the time course and corresponding to time points where there is no MI in the raw EEG voltage. This indicates that in those regions, even though observing the recorded voltage at a single time point does not reveal anything about the value of the stimulus feature the relationship between nearby time points is highly informative.

**Figure 13.**
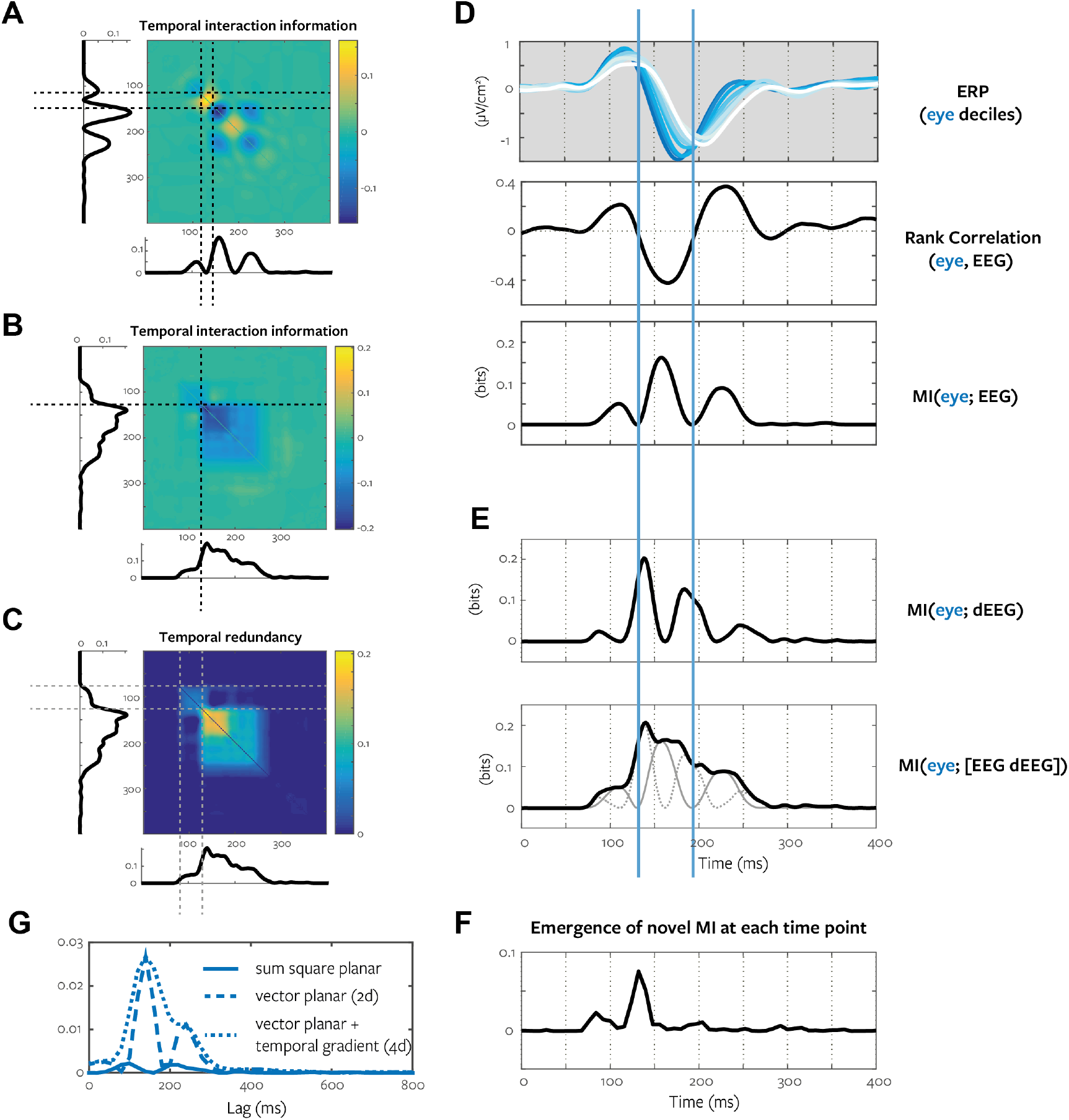
*Temporal interaction information reveals modulation of gradient*. **A.** Interaction information between EEG voltage at pairs of time points. Positive values correspond to synergy, negative values indicate redundancy. **B.** Interaction information between bivariate EEG voltage and temporal derivative at pairs of time points. **C.** As B, but only redundancy is shown. **D**. The mean ERP was calculated separately for each decile of the stimulus feature (white to blue increasing eye visibility). Spearman correlation and MI are calculated for the EEG voltage at each time point. **E.** The MI time course is calculated using the temporal derivative (upper) and a bivariate response consisting of the EEG voltage and temporal derivative at each time point (lower). This bivariate MI time course (black) is shown with the MI time courses of the constituent variables (EEG voltage, solid gray; temporal derivative, dotted gray lines). **F**. We down-sampled the data to 125 Hz, and calculated the new MI arriving at each time point (see text). **G**. Effect of including the temporal derivative with the 2d planar gradient response from Section 4.2, Figure 12.

The simplest quantification of the relationship between neighboring time points is the temporal gradient. To determine if this could account for the observed synergy we calculated the central difference temporal gradient of the EEG voltage for each trial and considered the MI in this response (Figure 13E, upper). Peaks in the gradient MI occur concurrently with zero points of the raw voltage MI. To incorporate the modulation of both response representations we combine them in a bivariate response consisting of the EEG voltage and the temporal gradient at each time point. We calculate the time course of MI about the stimulus feature in this bivariate response (Figure 13E, lower). Relating this to the voltage MI time course and the actual ERP modulation (Figure 13D) we can see that the zero points of the triple-peak MI profile result from zero-crossings where the sign of the correlation changes, due to the shape of the modulated bimodal ERP. However, by considering the conditional ERPs it is clear that these points fall within the time window where the overall shape of the evoked EEG response is modulated by the stimulus feature. We therefore take the view that considering the gradient together with the voltage (Figure 13E) provides a substantial advantage: these artifactual dips are smoothed out, providing a clearer picture of the time window over which the EEG signal is modulated by the changing stimulus.

We suggest that including the gradient of recorded neuroimaging signals could be a useful principle across a range of different analyses. For example, returning to the MEG data set under continuous speech stimulation (Section 4.2), including the temporal derivative of each planar gradient component (resulting in a 4d response) has the same effect of smoothing out the artifactual MI zero resulting from a change of sign in the effect (Figure 13G). Again, this gives a clearer, smoother picture of the range of delays over which the amplitude of the speech envelope modulates the MEG signal, which is made possible with the use of a multivariate statistic.

We can repeat the temporal interaction analysis on our bivariate responses (Figure 13B). This reveals that there is now little synergy, the main MI peak is mostly self-redundant, but the early part of the MI time-course appears to be independent from the main peak. To give a clearer picture we show the redundancy only (negative II; Figure 13C). A block structure is clearly apparent; the early MI appears to be independent from the bulk of the later MI (indicated with dashed lines). This suggests a functional differentiation between the initial P100, and the later N170: they appear to be modulated by the eye visibility in different ways and so possibly reflect different processing pathways. This would not be apparent from inspecting the ERP modulation (Figure 13D) or the MI time course (Figure 13E) alone.

Another application of the copula MI framework allows us to directly quantify the emergence of novel MI over time. For each time point, we calculate the MI about the stimulus feature available in the (bivariate) EEG response at that time point. We then subtract the MI that is redundant with that at the previous time point, leaving only the amount of new MI about the stimulus arriving at that time. Mathematically, this is equivalent to calculating the CMI *I* (eye; *R*_*t*_i__ | *R*_*t*_*i*_−1_). Figure 13F shows the result of this analysis. Two peaks of novel MI are clearly visible. In this analysis, for later time points the response at the time of the two peaks are also conditioned out to ensure only genuinely new MI is measured. The first peak corresponds to the early P100 modulation, the second to the stronger N170 modulation. This analysis corroborates the temporal interaction presented above, revealing that there appear to be two separate processes modulated by the stimulus feature – one beginning at 84 ms (P100), and one beginning at 132 ms (N170).

In summary, we have shown a few illustrative examples of the application of pairwise interaction information, focusing on interactions in the temporal domain with EEG data. We have shown how viewing pairwise interactions in terms of synergy and redundancy about a stimulus feature can provide useful insights – for example by revealing the importance of considering the temporal derivative when evaluating stimulus modulation of an evoked signal, and allowing us to directly quantify the emergence of new information over time. We emphasize that this approach is completely general and can be used across wide range of different responses (Figure 7A).

### 4.4. Bias and Mean-square-error of the GCMI estimator

We have so far focused primarily on the properties of the GCMI estimator when used for a permutation-based null-hypothesis significance test. This is for two reasons. First, the null-hypothesis statistical testing approach is widely used in neuroimaging and is perhaps the most likely application for most users of our new estimator. The performance in terms of sensitivity and specificity in comparison with existing statistical techniques is of crucial interest for such users (Sections 4.1, 4.2). Second, as described in Section 3, the GCMI estimator provides a lower-bound estimate to MI. This lower bound property complicates direct interpretation of the estimated MI quantities. In this section we explicitly address this issue.

Figure 14 shows estimated MI as a function of sample size for various systems and MI estimators. We first consider a bi-variate Gaussian system (1d stimulus, 1d response) with different levels of correlation. Figure 14A shows the expected value (mean) and variation (error bars show 25^th^ – 75^th^ percentiles) over 500 independent simulations, as a function of the number of samples (log scale). For Gaussian systems the true value can be calculated analytically and is indicated with a dashed line. For comparison we include binned methods, with 2 and 4 equi-populated bins for each signal with Miller-Madow bias correction applied (Miller, 1955). We also include the Kraskov-Stögbauer-Grassberger k-nearest-neighbor method as one of the most widely used continuous MI estimators (Khan et al., 2007; Kraskov et al., 2004; Lindner et al., 2011; Lizier, 2014). Across the range of sample sizes considered, the GCMI estimator has similar bias to the KSG estimator, but considerably lower variance, which results in systematically lower mean-square-error (lower panels). The 2-bin method has similar bias to GCMI; 4-bins suffers from larger bias. However, the MSE for these binned measures is substantially higher, because even in the large sample limit they systematically under-estimate the true continuous information (although the estimate gets closer with a higher number of bins). Figure 14B shows a similar simulation in a multivariate case – here a tri-variate Gaussian representing a uni-variate stimulus which modulates both components of a two-dimensional response. We observe similar relationships between the methods; GCMI has lower bias, lower variance and lower MSE than the KSG estimator. The binned methods suffer from increased bias and again under-estimate the continuous MI.

**Figure 14.**
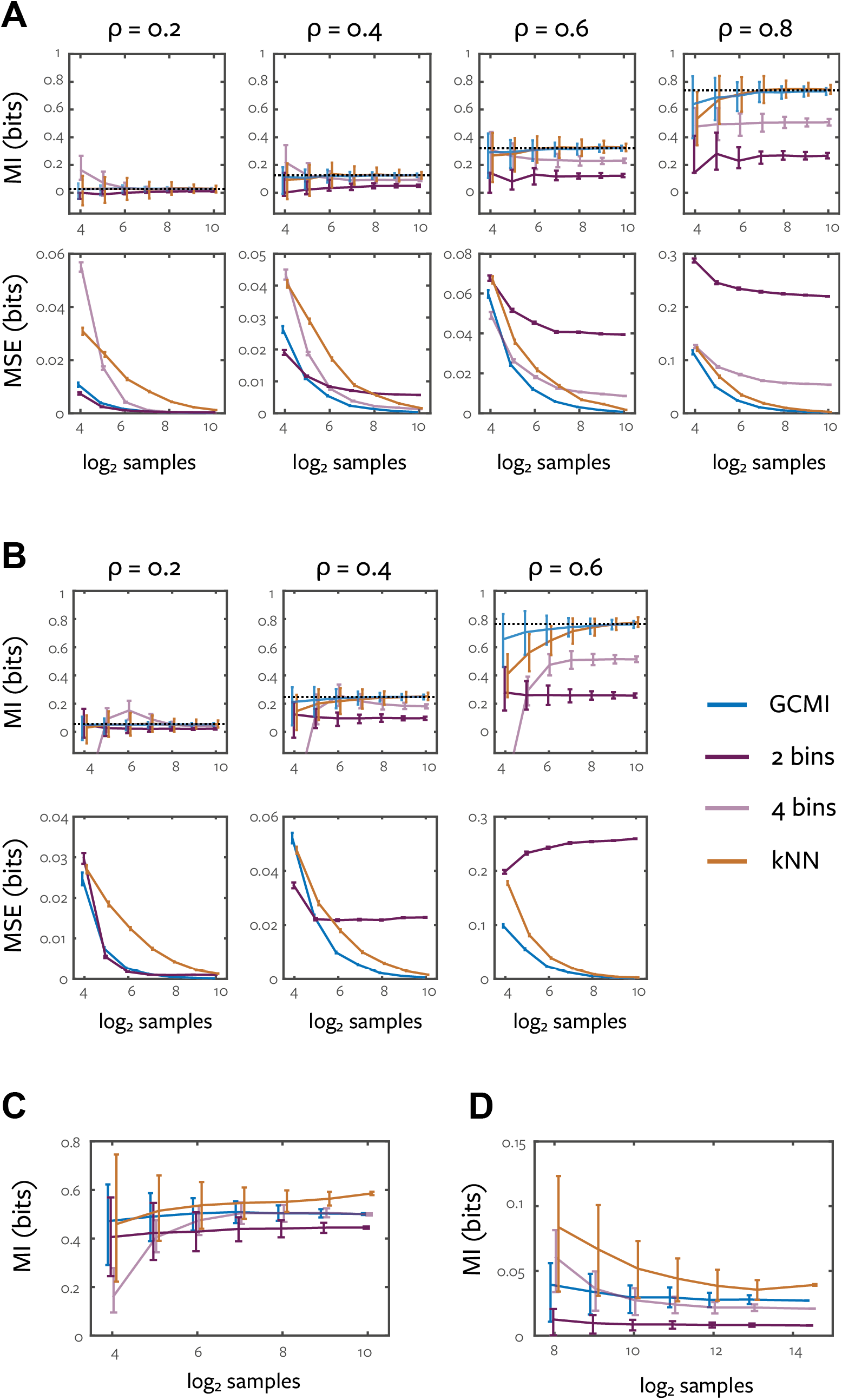
*Bias properties of the Gaussian-copula mutual information estimator*. **A.** Data were simulated from bivariate Gaussian distributions with 4 levels of correlation (0.2,0.4,0.6,0.8) and MI between the two variables was calculated with a range of methods. Upper panels show mean (error bars show 25^th^ – 75^th^ percentiles) over 500 simulations as a function of the number of samples (log scale) for 4 different MI estimators (see text): GCMI (blue), KSG nearest-neighbour estimator (k=3, orange), 2 and 4 equi-populated bins (dark purple, light purple respectively). Binned estimates are corrected with Miller-Madow bias correction. Lower panels show mean square error of the methods (error bars show s.e.m.) compared to the analytic ground truth value. **B.** The same simulation framework was applied to data sampled from a tri-variate Gaussian. One variable represented the stimulus and was correlated to a varying degree (0.2,0.4,0.6) with each of the response variables (which were themselves weakly correlated with r=0.1). **C.** 500 sub-samples of different sizes were drawn from the two-class event-related EEG data set described in Section 4.1. Plot shows mean (error bars show 25^th^ – 75^th^ percentiles). **D.** 500 sub-samples of 5s blocks were drawn from the continuous MEG data set described in Section 4.2. Plot shows mean (error bars show 25^th^ – 75^th^ percentile).

For these simulations, the dependence between the variables by construction does follow a Gaussian copula, hence the lower bound of the GCMI estimate is tight. With real data this is not necessarily the case. We performed similar analysis of the bias of the estimator with the experimental data presented in Sections 4.1 and 4.2. Bootstrap sampling (with replacement) is not suitable for use with the nearest neighbor based KSG estimator due to the effects of repeated data points on the nearest-neighbour calculation (Abadie and Imbens, 2008). We therefore sub-sample data sets without replacement. Figure 14C shows the results for the two-class event-related EEG dataset considered in Section 4.1 with the same channel as Figure 11B. Here we apply the Kozachenko and Leonenko nearest-neighbour entropy estimator (Kozachenko and Leonenko, 1987) to the class-conditional and unconditional data set and calculate MI following Eq. (3). Due to the combinatorial properties of subsampling from the 1078 trials, there is less variation in the largest data sample: the value there is close to that measured from the entire data set. The GCMI measure produces similar estimates to the 4-bin discrete method, with similar asymptotic value, but lower variance and much lower bias at small samples. The k-NN method does produce a slightly higher estimate suggesting that there maybe some non-Gaussian copula dependence in this data set. However, as shown in Section 4.1, the GCMI still provides an effective and sensitive statistical test when combined with permutation testing and the method of maximum statistics. Figure 14D shows a similar sub-sampled analysis for the continuous MEG data set, using the channel and optimal delay lag shown in Figure 12B. Here GCMI provides a higher estimate than either of the binned methods, with reduced bias (but similar variance). The KSG method appears to reach a slightly higher asymptotic value than the GCMI, but it is difficult to determine this without a larger data set. The KSG method seems to have a larger bias and variance here – we suspect this is due to the nearest-neighbour approach being more strongly affected by autocorrelation between nearby temporal samples.

In general the GCMI estimate may be systematically lower than the true MI value, even in the large sample limit. Standard techniques such as the bootstrap (Efron and Tibshirani, 1994) can be used to determine the sampling variability of the estimator, but such techniques cannot address the deviation of the GCMI estimate from the true MI. Any deviation of the empirical data copula from a Gaussian copula will lead to an underestimate of the MI, due to the maximum entropy property of the Gaussian copula. An extreme example in the uni-variate case is *y* = |*x*|+ε, with x a standard normal. In this case due to the symmetry in the empirical copula, the GCMI estimate will report 0 bits of information, while the true value can be arbitrarily large depending on the noise level (ε). This example suggests that for uni-variate variables the GCMI is sensitive to the same effects as a rank-correlation. Another likely source of mismatch with the Gaussian copula is the presence of tail-dependence in the data. For example, for t-distributed data, the Gaussian copula will have higher (negative valued) entropy than the true t-copula (which includes higher density tails), and therefore GCMI will be an underestimate, with the deviation increasing with correlation strength and decreasing with the t-distribution degrees of freedom.

While there are statistical tests for goodness-of-fit (GOF) of specific copulas (Genest et al., 2009b; Malevergne and Sornette, 2003) it is unclear how to directly relate any copula GOF test effect size to the tightness of the GCMI lower bound. For example, with multivariate responses there could be a strong deviation from the Gaussian-copula between the response variables, but in a way that does not affect the relationship between the stimulus and the multivariate response. The rejection of the hypothesis of a Gaussian copula does also not seem particularly useful, since with sufficient data that hypothesis could be rejected even when there is a very small difference between the GCMI and the true MI estimate.

Despite this, we propose the GCMI estimator is a useful practical tool, as a lower-bound MI estimate quantifying Gaussian-copula dependence, and particularly as an effect size for a flexible approach to permutation based statistical testing in a range of situations (Table 1). We have shown that with typical neuroimaging data it performs similarly to binned methods, but with generally better sampling properties. Binned methods similarly provide a lower-bound to the true continuous MI (see Figures 14A,B) but have nonetheless been extensively applied in practice to yield fruitful results (Section 1). GCMI is computationally much more efficient than the nearest-neighbour based method, with lower variance, and better sampling bias properties. As long as users keep in mind they are measuring only Gaussian-copula dependence (as they are with most existing classical statistics) the GCMI effect size provides a useful estimate of mutual information. As demonstrated (Section 4.1,4.2), while it may underestimate the true MI, it nonetheless has comparable sensitivity and specificity as conventional statistics when applied to mass-univariate (or mass-multivariate) permutation based inference in neuroimaging.

## 5. Discussion

Information theory provides a principled methodology for studying and quantifying statistical relationships between variables. As we have reviewed, the foundational quantities of information theory are entropy and mutual information. Here we have presented a novel approach to the practical estimation of these quantities, combining the statistical theory of copulas with the closed form solution for the entropy of Gaussian variables. This provides a computationally efficient and statistically robust lower bound estimate to MI with no specific assumptions on the marginal distribution of each variable. We have validated the use of this estimate as a statistical test within a neuroimaging context, considering both discrete and continuous experimental stimuli, and have shown that with 1D responses it performs as well as existing commonly used statistics. To accompany this article, we have released open-source code implementing the new methods for both Matlab and Python programming languages, together with tutorial examples covering the analyses presented here. The major advantage of our method over traditional statistical approaches is that it unifies a variety of applications (continuous, discrete and multidimensional variables) in a framework with a common effect size, and quantities like CMI and interaction information allow novel interpretations that are not available with other approaches.

The package implementing the approach in Matlab and Python is available at: https://github.com/robince/gcmi

Code for all simulations and figures is available at: <u>https://github.com/robince/sensorcop</u>

### Application to multi-dimensional spaces

A particular advantage of the proposed method is the ability to estimate MI and other information theoretic quantities on multi-dimensional spaces. We suggest that there are many situations in neuroimaging where multivariate responses are interesting, but difficult to address with existing statistical methods. Our examples included considering complex MEG spectra (as well as separating effects on phase and amplitude), 2D planar magnetic field gradients and considering raw signal values together with the instantaneous temporal derivative. Similarly, we could consider 3D magnetic fields arising from MEG source localization techniques, higher order temporal derivatives or features describing single-trial ERP features (peak and latency) (Hu et al., 2010). The multivariate performance is also important for calculating higher order information theoretic quantities such as conditional mutual information and interaction information. This is challenging with existing methods due to the curse of dimensionality, which either results in excessive data requirements (binned methods) or high computational complexity (continuous methods), even for modest numbers of dimensions (i.e. pairwise interaction information on 2D response variables as in the example of Section 4.3 requires estimation of entropy over a 5D space). While a more thorough analysis of the data requirements of the proposed method is an important area for future work, our experience to date is that with amounts of data that can reasonably be collected in a suitably designed neuroimaging experiment (i.e. hundreds of trials) spaces of dimension 5-10 can reliably be addressed, although of course this depends on the strength of the underlying effects. For much greater numbers of dimensions estimation of the required covariance matrices becomes problematic, even given the improved robustness resulting from the copula rank transformation.

The application of regularization through Bayesian priors or other means (Engemann and Gramfort, 2015) might provide a way to extend the measure to even higher dimensional spaces. Alternatively, we propose that one way to extend our estimator to very high-dimensional spaces is to combine it with decoding approaches based on supervised learning algorithms. For example, by first using a decoding algorithm (e.g. a linear discriminant) as a dimensionality reduction step, we can calculate the MI of the low-dimensional predictor signal, within a cross-validation framework. We suggest that MI has some advantages as a statistic to evaluate the performance of a decoder (Quian Quiroga and Panzeri, 2009) compared to commonly used measures (such as mean performance, area under ROC curve, etc.). Again it uses a common scale, provides the ability to relate MI in different signals (e.g. between EEG sensor array linear discriminant output and single-trial fMRI voxel beta activations, see below) and allows us to condition out correlated features. Other approaches to estimating MI in higher dimensional response spaces include extensions to the nearest-neighbour method with specifically chosen distance measures that preserve the appropriate structure of the high-dimensional space. For example, in fMRI a distance based on correlation between voxel time courses can be used to estimate MI between a statistical parameter map and a high-dimensional whole brain validation data set (Afshin-Pour et al., 2011).

### Second level analyses on MI values

MI provides a high contrast statistic that can be used as input for second level analyses. MI reveals a functional property of the system that might change with different experimental conditions, for example with the rhythmic structure of speech stimuli (Kayser et al., 2015) or spatial attention (Guggenmos et al., 2015; Saproo and Serences, 2010). Functional connectivity, measured with directed information, has been shown to be affected by the intelligibility of speech (Park et al., 2015). When considering MI computed in different experimental conditions accurate bias correction is important because bias may not be equal in each condition (for example due to differing numbers of samples, or different degrees of signal autocorrelation). Mass-univariate MI calculations can also provide a rich and descriptive input for subsequent dimensionality reduction. For example Non-negative Matrix Factorization (NMF) (Lee and Seung, 1999) is a dimensionality reduction technique that is well matched for application to MI results (since they are non-negative, and the high signal-to-noise contrast of MI complements the mean squared-error objective of the NMF algorithm). This can be used to extract task relevant features from a high-dimensional naturalistic stimulus (Ince et al., 2015); the same approach could also be applied to reduce the dimensionality of a high-dimensional neuroimaging response (e.g. an EEG sensor array) into specific spatio-temporal task or stimulus MI components.

### Quantifying pairwise interactions

We believe that understanding brain function from brain activity requires a focus on the particular information processing functions performed under different tasks and conditions (Kriegeskorte et al., 2006; Schyns et al., 2009). To fully exploit the potential of this information processing perspective requires methods that not only identify the systematic modulations of brain responses, but the relationship between such modulations, or representations, across different times, regions and signals (Figure 7A). The use of such approaches within neuroimaging is growing, with development and application of techniques such as Representation Similarity Analysis (RSA) (Kriegeskorte and Kievit, 2013), and the use of supervised classification algorithms together with cross-validation (Hastie et al., 2001), often referred to as a decoding (Haxby et al., 2014; King and Dehaene, 2014; Quian Quiroga and Panzeri, 2009). Our new MI estimator allows us to address these issues within the unified framework of information theory. Particular advantages include the common, meaningful scale that allows direct comparison of redundancy with the MI in different signals, experiments, or behavior. With information theory we can normalize redundancy as a percentage, which provides a more intuitive measure for the degree of overlap than is available with other methods, and we can perform all analyses conditioning out multiple correlated stimulus features if necessary. Detailed comparison of our information theoretic framework with methods such as RSA will be the subject of future work.

Our novel estimator can be applied to calculate measures of functional connectivity such as directed information (transfer entropy). In combination with the information perspective described above, by adding the concept of redundancy within the Granger causal framework we have developed a measure of functional connectivity that quantifies the communication of specific information content (Ince et al., 2015). The copula MI estimate is crucial to allow practical computation of this measure, which requires conditioning directed information on the particular stimulus features considered. This measure allows for dynamic network analyses that are based not on general relationships between areas, but on communication of specific information about the stimulus or task.

### Broader applications

We have focused here on application to M/EEG data, but we emphasize that the copula MI estimator can be applied to any signal, facilitating the comparative study of neural information coding across experimental methodologies and scales of brain measurement (Panzeri et al., 2015). For example, it could be applied to single trial fMRI GLM beta activations directly, or in combination with a multi-voxel decoding approach as described above. The common scale allows direct comparison of the strength of the modulation between different neuroimaging responses, and interaction information opens up the promising possibility of directly relating the information content in different signals (Cichy et al., 2014). For example, calculating the redundancy over time between MI in a (multivariate) EEG response and individual voxel single-trial beta activations, would allow mapping the spatial region that is redundant with the EEG at each time point.

Similarly, while we have focused here on neuroimaging, MI has broad applicability as a general statistical framework. It can be used for analyzing behavioral data, where many of the properties we have highlighted could be useful (e.g. CMI, interactions). It has been used for feature selection in general classification problems (Lefakis and Fleuret, 2014; Peng et al., 2005; Torkkola, 2003) and we hope our estimator would also provide practical advantages in many such applications. We further suggest that the copula normalization could be used as a general preprocessing step that would convert any covariance based statistic or algorithm into a robust rank-based version (e.g. common spatial patterns, canonical correlation analysis, linear/quadratic discriminant analysis).

### Conclusion

We have presented a novel approach to estimate MI and associated quantities. This provides a general, computationally efficient, flexible, robust, multivariate statistical framework based on information theory. This framework provides effect sizes on a common meaningful scale and allows for unified treatment of discrete and continuous variables. Beyond measuring the strength of direct, possibly multivariate, relationships, quantities like CMI and interaction information have the potential to provide transformative interpretations of neuroimaging data, for example by relating information content in different brain responses. This framework allows investigators to take full advantage of the properties of each neuroimaging signal and their experimental designs to develop a better understanding of the information processing functions of brain networks.

## Acknowledgements

We thank Arno Onken, Daniel Chicharro and Stefano Panzeri for useful discussions. JG and PGS are supported by the Wellcome Trust [098433, 107802]. GAR, PGS, BG and CK are supported by the UK Biotechnology and Biological Sciences Research Council (BBSRC) [BB/J018929/1, BB/D01400X1, BB/M009742/1, BB/L027534/1]. CK is supported by the European Research Council (ERC-2014-CoG; grant No 646657).

The authors report no financial conflict of interest.

